# A novel interaction between the 5’ untranslated region of the Chikungunya virus genome and Musashi RNA binding protein is essential for efficient virus genome replication

**DOI:** 10.1101/2023.03.29.534719

**Authors:** Kaiwen Sun, Francesca Appadoo, Yuqian Liu, Marietta Müller, Catriona Macfarlane, Mark Harris, Andrew Tuplin

## Abstract

Chikungunya virus (CHIKV) is a re-emerging, pathogenic alphavirus that is transmitted to humans by *Aedes spp*. mosquitoes—causing fever and debilitating joint pain, with frequent long-term health implications and high morbidity. The CHIKV replication cycle is poorly understood and specific antiviral therapeutics are lacking. In the current study, we identify host cell Musashi RNA binding protein-2 (MSI-2) as a proviral factor. MSI-2 depletion and small molecule inhibition assays demonstrated that MSI-2 is required for efficient CHIKV genome replication. Depletion of both MSI-2 and MSI-1 homologues was found to synergistically inhibit CHIKV replication, suggesting redundancy in their proviral function. Electromobility shift assay (EMSA) competition studies demonstrated that MSI-2 interacts specifically with an RNA binding motif within the 5’ untranslated region (5’UTR) of CHIKV and reverse genetic analysis showed that mutation of the binding motif inhibited genome replication and blocked rescue of mutant virus. For the first time, this study identifies the proviral role of MSI RNA binding proteins in the replication of the CHIKV genome, providing important new insight into mechanisms controlling replication of this significant human pathogen and the potential of a novel therapeutic target.

## INTRODUCTION

Chikungunya virus (CHIKV) is transmitted by *Aedes spp.* mosquitos and is an alphavirus of the *Togaviridae* family, which includes other medically relevant alphaviruses such as Semliki Forest virus (SFV), Venezuelan equine encephalitis virus (VEEV), and Sindbis virus (SINV) (1,2). CHIKV was first identified in Tanzania in 1952 and symptoms typically include acute febrile symptoms, myalgia, rash and severe arthralgic joint pain, which may persist for months or years (3). CHIKV recently caused epidemic outbreaks across regions of Asia, Africa, the Americas, the Middle East and Southern Europe (3). To date, three phylogenetically distinct lineages of CHIKV have been identified, namely the West African, Asian and the East Central Southern African (ECSA) lineages (4). There remains no clinically approved specific antiviral therapy, due in part to a lack of detailed understanding of the CHIKV replication cycle and its interaction with host cell factors. The first attenuated live CHIKV vaccine (VLA1553) received U.S. FDA approval in November 2023 (5).

CHIKV is an enveloped, positive-sense single-stranded RNA virus with an ∼11.8 Kb genome, containing two open reading frames (ORFs) flanked by 5’ and 3’ untranslated regions (UTRs) (**Fig. 1A**). The 5’ UTR is ∼ 76 nts in length and is capped by 5’ type-0 N-7-methylguanosine. The upstream ORF (ORF-1) encodes the viral non-structural proteins 1-4 (nsP1-4), which are translated directly from the genomic RNA as a single polyprotein that is subsequently proteolytically cleaved into the four mature proteins (6). Through analogy with other alphaviruses, proteolytic cleavage of nsP1-4 in *cis* by nsP2 releases nsP4 which, as the RNA-dependent RNA polymerase (RdRp), initiates synthesis of the minus-strand intermediate RNA. Subsequent proteolytic cleavage of the remaining nsP123 polyprotein initiates replication of genomic and sub-genomic (29S) RNAs from the minus-strand template (7). The downstream ORF (ORF-2) encodes the structural polyprotein that is processed into the capsid and envelope proteins, E3, E2, 6K, and E1.

**Figure 1.**
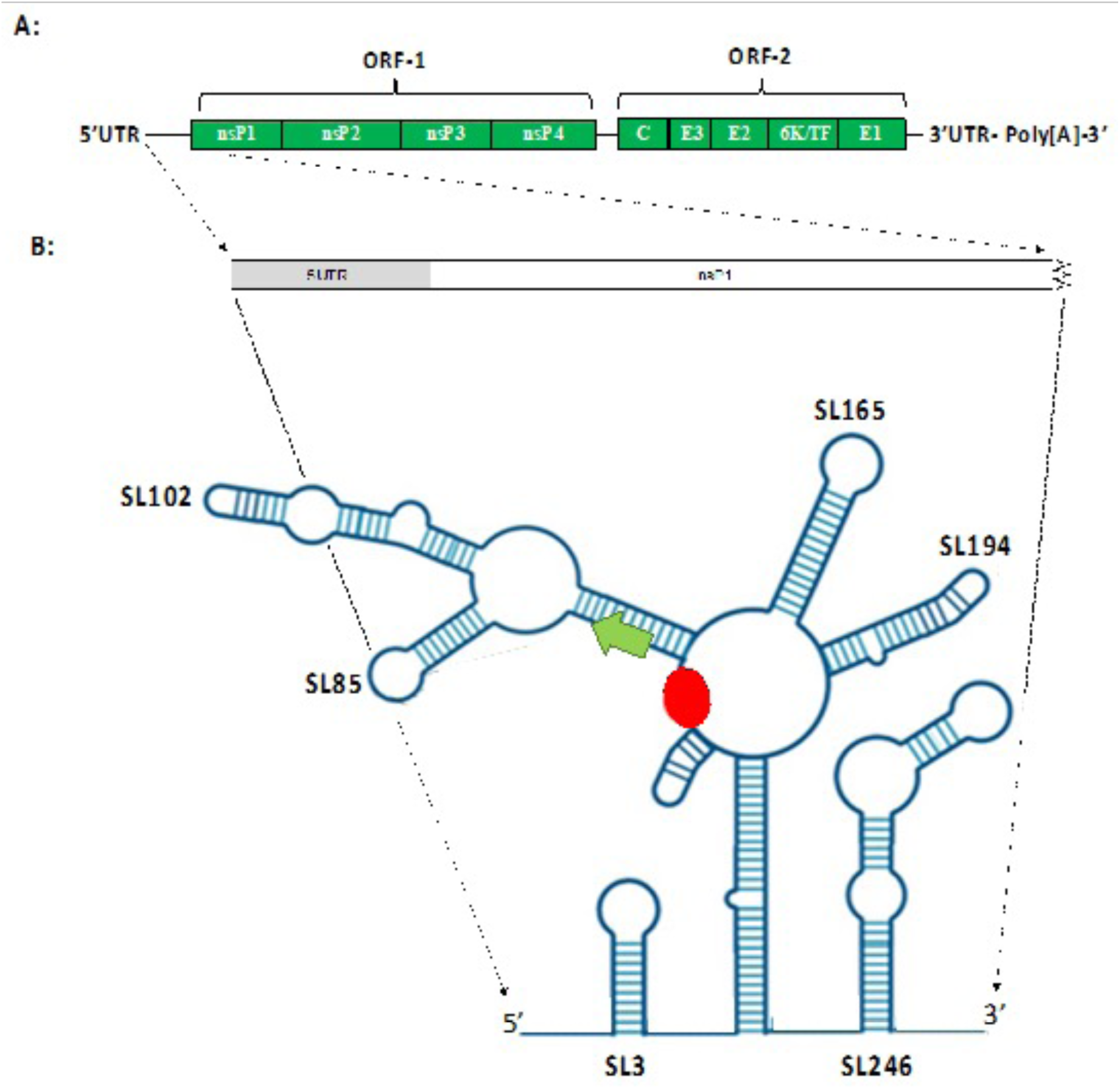
**A)** Schematic representation of CHIKV genome organisation **B)** Schematic representation of CHIKV RNA structures within the 5’UTR and adjacent ORF-1 region of the CHIKV genome (Kendall et al., 2019). RNA replication elements SL3, SL47, SL88, SL102, SL165, SL194 and SL246 are labelled in black type. The ORF-1 AUG start codon is labelled by a green arrow and the putative MSI binding site by a red oval.

While negative strand replication is initiated at the 3’ end of the virus genome, as with many positive single stranded RNA viruses, regulatory elements and interactions within the 5’ end of the CHIKV genome are required for its initiation and regulation (8). We and others have previously demonstrated that such regulatory elements include functional RNA secondary structures and higher order interactions within the CHIKV 5’UTR and adjacent upstream region of ORF-1 (Fig 1). It has been speculated that these RNA elements may regulate template specificity and temporal control of switching, between CHIKV translation and genome replication (9,10).

In a previously published study, we demonstrated by SHAPE mapping and reverse genetic analysis that the CHIKV 5’UTR and adjacent ORF-1 coding region is highly structured (10). However, the study also highlighted a single-stranded region (nts 63-69) exhibiting a high degree of nucleotide conservation that was located immediately between two RNA structures (Supplementary data 1), which we demonstrated were essential for initiation of CHIKV genome replication (Fig 1B).

Interestingly, *in silico* analysis noted that nts _63_AUUAAU_68_ were closely homologous to the Musashi RNA binding protein (MSI) consensus binding sequence ((G/A)U_1-3_AGU). MSI are highly conserved RNA binding proteins, containing two highly conserved tandem RNA recognition motifs (RNP-1 and RNP-2), that interact with RNA via the same consensus binding motif. Two MSI homologues have been identified, MSI-1 and MSI-2, that share over 90% homology in their RNA-binding domains and a high degree of functional complementarity and redundancy (11,12). MSI have key roles in post-transcriptional regulation of genes involved in development, cell cycle regulation and maintenance of adult neural stem/progenitor cells (13). A recent study demonstrated that MSI-1 promotes Zika Virus (ZIKV) genome replication in neurons, via interaction with the consensus binding site within the viral 3’UTR (14).

In the current study, we demonstrate through a range of infectious clone and sub-genomic replicon reporter assays that MSI-2 is required for efficient CHIKV genome replication. A small molecule inhibitor of MSI-1 and MSI-2 reduced infectious virus production by inhibition of CHIKV genome replication. Similarly, MSI-2 silencing by shRNA and siRNA inhibited both infectious virus production and genome replication, while co-silencing of both MSI-2 and MSI-1 exhibited a synergistic effect on inhibition of CHIKV replication. Reverse genetic analysis, in which the putative MSI binding site within the CHIKV 5’UTR was mutated, completely prevented rescue of mutant virus and inhibited CHIKV genome replication. Biochemical MSI-2 binding analysis by competition electromobility shift assay (EMSA) demonstrated that recombinantly expressed MSI-2 specifically interacts with the 5’ region of the CHIKV genome and that this interaction is inhibited by mutation of the putative MSI 5’UTR binding site. These findings demonstrate for the first time that interaction between MSI and a binding site within the 5’UTR are required for replication of the CHIKV genome.

## Materials and Methods

### Cell culture

Human Rhabdomyosarcoma (RD), Human hepatoma (Huh7), Baby Hamster Kidney (BHK) and Human Embryonic Kidney (HEK) 293 were grown in Dulbecco’s modified eagle’s medium (DMEM; Sigma) containing 10% foetal bovine serum (FBS; PAA), 1x penicillin-streptomycin (Sigma), 25mM HEPES in 0.85% NaCl (Lonza) and 1% non-essential amino acids mixture (NEAA; Lonza). Cells were harvested using trypsin/EDTA (Sigma) and maintained at 37°C in 5% CO_2_.

### CHIKV cDNA plasmid

The infectious CHIKV (ICRES), CHIKV Dual-luciferase sub-genomic replicon (CHIKV-SGR) and *trans-*complementation (pCHIKV-nsP1234 and pCHIK-Fluc/Gluc) assay were derived from the CHIKV ECSA strain, isolate LR2006 OPY1 (accession number DQ443544)(15). In CHIKV-SGR the second ORF was replaced by a firefly luciferase gene and a *Renilla* luciferase gene was fused within nsP3 (16). The *trans-* complementation assay utilised a CMV codon optimised plasmid to supply the CHIKV replicase in *trans* (pCHIKV-nsP1234) and a Pol II expressed reporter plasmid (pCHIKV-FLuc/GLuc), in which the majority of ORF-1 (downstream of nt 320) was replaced by a firefly luciferase gene and all of ORF-2 by a Gaussia Luciferase gene (17). Plasmid cDNA was purified using GeneJET Plasmid Maxiprep kits (Thermo Fisher Scientific) according to the manufacturer’s instructions.

### *In vitro* RNA transcription

To generate CHIKV-SGR and infectious CHIKV RNA, 2 μg of cDNA was linearised with Not-I HF and used as a template for transcription of 5ʹ [m7G(5ʹ)ppp(5ʹ)G] capped (m7G capped) RNA from an upstream SP6 promoter, using an SP6 mMessage mMachine kit, according to the manufacturer’s instructions (Life Technologies). Uncapped CHIKV 1–337 RNA, used 1 μg of CHIKV 1–337 PCR DNA as a template for *in vitro* transcription using the SP6-Scribe™ standard RNA IVT kit, according to the manufacturer’s instructions (Lucigen). In all cases, following DNase I treatment, RNA was purified by LiCl precipitation and analysed by denaturing agarose gel electrophoresis.

### Virus production

1 × 10^6^ BHK cells were trypsinised and resuspended in 400 μl ice-cold DEPC-PBS. Cells were then electroporated with 1 μg 5ʹ-capped CHIKV ICRES *in vitro* transcribed RNA in a 4 mm electro-cuvette, with a single square wave pulse at 260 V for 25 ms using a Bio-Rad electroporator, before seeding into a T175 flask in 20 ml DMEM. After 24 h, supernatant was aspirated and virus titre measured by plaque assay on BHK cells.

### CHIKV quantification by plaque assay

BHK cells were seeded at 1 × 10^5^ cells per well in 12-well plates and maintained overnight in 1 ml DMEM. The following day monolayers were washed with PBS, infected with 10-fold serial dilutions of CHIKV transfection supernatant and maintained at 37°C. At 1 hour post infection (h.p.i.) monolayers were washed with PBS and covered with a 0.8% methylcellulose DMEM P/S overlay. Monolayers were fixed and stained at 48 h.p.i. (5% paraformaldehyde and 0.25% crystal violet respectively), plaques counted and virus titres expressed in plaque-forming units per ml (PFU/ml).

### Ro 08-2750 cell viability assay

RD cells were seeded in 96-well plate at 8×10^4^ cells/well and maintained for 24 h. Monolayers were then treated with increasing doses (0, 0.5, 1, 3, 5, 7, 10, 20 μM) of the MSI small molecule inhibitor Ro 08-2750 (TOCRIS) dissolved in DMSO. After 24 h media/inhibitor was aspirated and replaced with 20 μl of 5 mg/mL 3-(4,5-dimethylthiazol-2-yl)-2,5-diphenyltetrazolium bromide (MTT) (Sigma) and incubated at 37°C in 5% CO_2_ for 3 h. After incubation, MTT solution was replaced with 100 μl DMSO and the plate shaken at 60 rpm for 5 min. Absorbance at 570 nm was determined using an Infinite F50 microplate reader (Tecan) and expressed as a percentage of DMSO control cells. 5 μM Ro 08-2750 was estimated to be the maximum non-toxic dose in in Huh7 cells and was used for all further assays.

### Infectious CHIKV Ro 08-2750 inhibition assays

RD cells were seeded in 12-well plates at 1×10^5^ cells/well and incubated overnight in the presence of Ro 08-2750. Monolayers were then infected with CHIKV at MOI=0.1 and adsorbed to the cells for 1 h at 37°C before aspirating and maintaining in the presence of Ro 08-2750 for 24 h. Supernatants was then collected and infectious virus production measured by plaque assay.

### siRNA depletion of MSI-1 and MSi-2

RD or Huh7 cells were seeded at 1×10^5^ cell/well in 12-well plates in antibiotic-free medium. After 24 h, cells were washed with PBS and incubated with 1x Opti-MEM + GlutaMAX (Gibco) for 20 min at 37°C/5% CO_2_. For each well, 50 pmol MSI-1 (sc-106836; Santa Cruz) and/or MSI-2 siRNA (sc-75834; Santa Cruz) were mixed with 100 μl Opti-MEM and incubated at room temperature for 1min. No siRNAs were added to mock samples and 50 pmol of scrambled siRNA (SI03650318; QIAGEN) was used as a negative control. In parallel, 3 μl Lipofectamine RNAiMAX (Invitrogen) was mixed with 100 μl Opti-MEM and incubated at room temperature for 1 min. The siRNA and Lipofectamine RNAiMAX were mixed and incubated at room temperature for 5 min before adding to the cells. After 24 h, cells were either lysed to confirm MSI-1 or MSi-2 depletion by western blot or used for subsequent CHIKV infection assays.

### shRNA depletion of MSI-1 and MSi-2

Human embryonic kidney 293 (HEK 293T) cells were plated in antibiotic-free DMEM in 6-well plates and maintained until confluency reached ∼80%. For each transfection, in a single tube 300 µL OptiMEM was mixed with 1 μg packaging plasmid p8.91, 1 μg envelope plasmid pMDG encoding VSV-G protein and 1.5 µg lentiviral vector plasmid encoding shRNA to MSI-2 (sc-75834-SH; Santa Cruz Biotechnology). A lentiviral vector plasmid encoding scrambled shRNA sequences with no cellular targets was used as the negative control (sc-108060; Santa Cruz Biotechnology). In another tube, 300 µL of OptiMEM was mixed with 5 µL lipofectamine 2000 (Invitrogen). Both tubes were gently mixed by flicking, incubated at room temperature for 5 min, mixed together and incubated for 20 min at room temperature. The antibiotic-free DMEM was aspirated and the monolayers washed once with PBS. 800 µL Opti-MEM was added to each well before dropwise addition of the lentiviral plasmids/shRNA mixture. After maintenance at 37°C for 6 h the media was changed to antibiotic-free DMEM. 48 hpt the lentivirus containing supernatant was harvested and filtered through a 0.45 µm filter.

RD cells were seeded at 1×10^5^ cells/well the day prior to transduction. 1 mL of the lentivirus supernatant and polybrene (MERCK) were added to each well and incubated for 6 h before aspirating and replacing with antibiotic free DMEM. After 72 h media was replaced with DMEM containing 2.5 μg/ml puromycin, in which the cells were then maintained. The efficiency of the shRNA MSI-2 depletion was confirmed by western blot.

### Strand-specific Quantification of CHIKV RNA

Total RNA was extracted from infected cells using TRI Reagent Solution (Applied Biosystems) according to the manufacturer’s instructions. Strand-specific RT-qPCR (ssRT-qPCR) was performed as previously described (18). Briefly, 500 ng of RNA was reverse-transcribed with gene specific primers (Supplementary data 2) using the SCRIPT cDNA Synthesis Kit (Jena Bioscience) according to the manufacturer’s protocol. 100 ng of strand-specific cDNA was used as template for the quantitative PCR performed with the qPCRBIO SyGreen Blue Mix Lo-ROX (PCR Biosystems), with gene specific primers amplifying a 94 bp region of the CHIKV nsP1-encoding sequence using the following PCR program: 95°C for 2 mins, 40 x (95°C for 5 sec, 60°C for 30 sec), dissociation curve 60°C-95°C as pre-defined by the Mx3005P thermal cycler (Agilent technologies). *In vitro* transcribed CHIKV ICRES RNA was reverse transcribed and a cDNA dilution series employed as a standard to quantify copy numbers in the respective samples.

### CHIKV Sub-genomic replicon assays

For analysis of Ro 08-2750 inhibition, RD cells were seeded in 24 well plates at 5 × 10^4^ cells per well and maintained overnight in the presence of Ro 08-2750. Monolayers were washed once in PBS before addition of 400 μl opti-Mem reduced-serum media and 100 μl transfection media. Transfection media was prepared according to the manufacturer’s instructions, using 1 μl Lipofectamine 2000 (Invitrogen), 250 ng of CHIKV-SGR RNA and appropriate concentrations of Ro 08-2750, before being made up to 100 μl using opti-Mem. Monolayers were maintained for 6 hpt before the media was aspirated and replaced with complete DMEM/Ro 08-2750. At 8 and 24 hpt monolayers were lysed with 100 μl 1 x passive lysis buffer according to the manufacturer’s instructions (Promega), stored at −80°C and analysed using Dual-luciferase substrate (Promega) in a FLUOstar Optima luminometer (BMG labTech). For shRNA MSI-2 depleted cell lines the same method was followed, with the exclusion of Ro 08-2750.

### *Trans-*complementation Assay

The *trans*-complementation assay was performed as previously described (19). Briefly, for analysis of Ro 08-2750 inhibition, RD cells were seeded in 12 well plates at 5 × 10^4^ cells per well and maintained overnight in the presence of Ro 08-2750. Monolayers were then co-transfected with 1 µg each of the pCHIKV-nsP1234 and pCHIKV-FLuc/GLuc using Lipofectamine 2000, as described previously, and assayed at 8 and 24 hpt. For shRNA MSI-2 depleted cell lines the same method was followed, with the exclusion of Ro 08-2750.

### Expression and purification of recombinant MSI-2 RNA binding domains

The MSI-2 RNA binding domain expressing plasmid pET-22HT-MSI-2 (amino acid 8-193 and referred to hereafter as MSI-2 8-193) was a kind gift from Prof S. Ryder (Addgene plasmid # 60356; http://n2t.net/addgene:60356; RRID: Addgene_60356) (20). The plasmid was transformed into BL21 (DE3) competent cells following the manufacturer’s protocol (NEB), a single colony was inoculated into 10 mL LB ampicillin (10 mg/ml) and incubated overnight at 37°C before inoculating 1000 ml 1 LB (ampicillin (10 mg/ml) and adding 100 μM Isopropyl β-D-1-thiogalactopyranoside (IPTG; Thermo Fisher Scientific) when the optical density reached 0.8. The culture was then incubated overnight at 18°C in an orbital shaker before centrifugation to pellet the bacteria. The pellet was resuspended in lysis buffer (20U/mL DNase I, 0.6U/mL RNase A, 1mg/mL lysozyme and 1x protease inhibitor) and lysed on ice for 30 min. Following sonication, the suspension was centrifuged twice at 2000 xg for 1 hour at 4°C. The was filtered through a 0.45 μm filter and His-MSI-2 was purified with HisTrap^TM^ FF column (Cytiva) using the Econo Gradient Pump (BIORAD) according to the manufacturer’s protocol and overnight dialysis. The purified protein was quantified and the purity and identity assayed by Coomassie SDS-PAGE and western blot (Supplementary data 3).

After desalting using PD-10 desalting columns (Cytiva), His-MSI-2 protein was further purified by ion exchange chromatography. Following the manufacturer’s instructions, the column was equilibrated with wash buffer (50 mM MES, 10 mM NaCl, pH 5.6, degassed) and the protein eluted in the same buffer. Ion exchange chromatography was performed using HiTrap SP HP cation exchange chromatography column (Cytiva) following the manufacturer’s instructions and eluted in degassed 50mM MES, 1M NaCl, pH 5.6. The eluted protein fractions were analysed by Coomassie SDS-PAGE and western blot (Supplementary data 3).

### 32P RNA end labelling

*In vitro* transcribed CHIKV RNA (nt’s 1-337) was 5’ dephosphorylated using Quick CIP according to the manufacturer’s instructions (NEB) before purification using an RNA Clean & Concentrator column (Zymo Research). 20 pmoles of dephosphorylated RNA was combined with 2 μl 10 x T4 Polynucleotide Kinase buffer (NEB), 3 μl ATP(y-^32^P) 10 mCi/ml, 1 μl T4 Polynucleotide Kinase (NEB) and nuclease free H_2_O to a final volume of 20 μl, incubated at 37°C for 60 min, 65°C for 20 min, purified using an RNA Clean & Concentrator Kit (Zymo Research) and resuspended in RNase-free H_2_O.

### Electromobility shift assay (EMSA)

RNA was incubated at 95°C for 2 min and on ice for 2 min before 3.3 x RNA folding buffer (100 mM HEPES pH 8.0, 100 mM NaCl and 10 mM MgCl2) and RNasin Plus RNase Inhibitor (Promega) was added to a final volume of 10 μl and incubated for 20 min at 37°C. MSI-2 protein was combined with 3.3 x RNA folding buffer, 0.5 x TE, 100% glycerol, RNasin Plus RNase Inhibitor (Promega) and 5 µg yeast tRNA as non-specific competitor and incubated at 37°C for 2 min before adding to the RNA mixture and further incubation at 37°C for 15 min. Finally, samples were analysed by native PAGE gel electrophoresis at 135 V for ∼2 h. The gel was fixed for 30 min and dried with gel dryer (BIORAD), exposed onto Hyperfilm™ ECL™ (Merck) and visualised using a Xograph Film Processor. Band shifts in the competition EMSAs were quantified by measuring the density of the RNA/Protein complex band (as indicated by square brackets) normalized to the density of the whole lane and expressed as % of the normalized RNA/Protein complex band in lane 1:0.

### Western blotting

Following infection and incubation, monolayers were lysed in IP lysis buffer (Promega) and incubated at room temperature for 30 min. Protein concentration was quantified using a Pierce™ BCA Protein Assay Kit (Thermo scientific) according to the manufacturer’s instructions. Equal amounts of protein lysate were analysed by SDS-PAGE. Protein was transferred onto an Immobilon-FL PVFD transfer membrane (MERCK) using a TE77X semi-dry transfer (Hoefer) at 15 V for 60 min. Membranes were blocked using diluted Odyssey® Blocking Buffer in PBS (LI-COR) for 30 min and probed with primary antibodies against β-actin (1:10,000, mouse monoclonal, Sigma-Aldrich A1978), MSI-2 (1:1000 rabbit monoclonal, Abcam ab76148) and MSI-1 (1:5000, Abcam ab21628) in diluted Odyssey® Blocking Buffer in PBS (LI-COR) overnight at 4°C. After overnight incubation, primary antibody was removed and membranes washed 3 times using PBS. Membranes were stained with secondary antibodies (IRDye® 800CW Donkey anti-Mouse; IRDye® 680LT Donkey anti-Rabbit; Li-Cor) for 1 h at room temperature, washed 3 times using 1 x PBS, dried and then imaged using an Odyssey® Fc Imaging System (Li-Cor).

### Statistical analysis

Statistical analysis was carried out using two-tailed Students Test comparisons on GraphPad Prism version 8.4.0. *P* values of ≤0.05 (*), ≤0.01 (**), ≤0.001 (***) were used to represent degrees of significance between each drug treatment/silencing/mutant to wild-type assay.

## RESULTS

### MSI-2 binds specifically to the predicted _63_AUUAAU_68_ 5’UTR MSI binding site in the CHIKV 5’UTR

In order to examine the potential for direct interaction between the RNA binding domains of MSI-2 and nucleotides _63_AUUAAU_68_ within the CHIKV 5’UTR, which had close sequence homology to the (G/A)U_1-3_AGU MSI consensus nucleotide binding motif, we used native EMSA (21). _63_AUUAAU_68_ is located in a single-stranded region of the 5’ UTR, 9 nts upstream of the AUG start codon and between two RNA structures that are essential for CHIKV genome replication (Fig 1B)(10). The first 330nts of the CHIKV genome (RNA-330), representing the 5’UTR and adjacent upstream region of ORF1, was *in vitro* transcribed, 5’ end radiolabelled with ATP-[γ-32P] and incubated at 37°C -conditions under which correct folding of the functional RNA-330 structure was previously validated (10). Following incubation with increasing concentrations of recombinantly expressed and purified MSI-2 8-193 in the presence of unlabelled tRNA, reaction products were separated by native PAGE and analysed by autoradiography (Fig 2A). Relative to unbound RNA-330, the presence of MSI-2 8-193 retarded RNA-330 migration during native PAGE, consistent with the formation of an RNA-330/MSI-2 8-193 complex, the increasing size of the retarded band potentially indicating MSI-2 multimerization. In order to investigate the specificity of the observed interaction, a fixed molar ratio of RNA-330^γ-32P^ and MSI-2 8-193 was incubated with increasing concentrations of competitor unlabelled RNA-330 (Fig 2B). Increasing concentrations of unlabelled RNA-330 reduced the formation of the RNA-330^γ-32P^/MSI-2 8-193 complex, with a corresponding increase in unbound RNA-330^γ-32P^.

**Figure 2).**
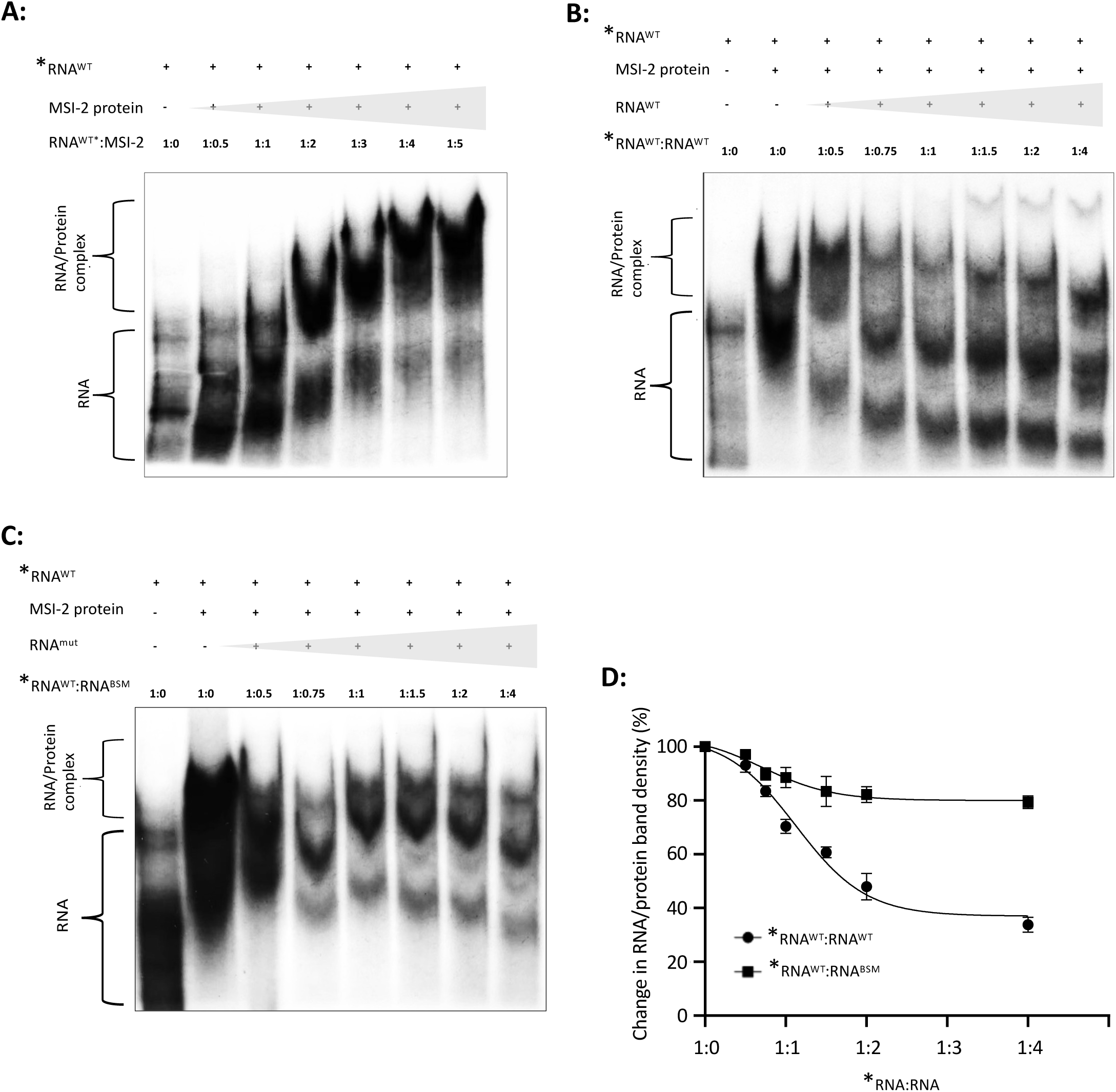
Native EMSAs between *in vitro* transcribed P^32^ 5’ radiolabeled CHIKV RNA nts 1-330 (*RNA^WT^) and recombinantly expressed MSI-2 demonstrated an RNA/protein interaction that was outcompeted by increasing concentrations of equivalent unlabeled (RNA^WT^) but less efficiently by the same RNA incorporating BSM mutation _63_AUUAAU_68_ >_63_CAACUU_68_ (RNA^BSM^). **A)** Increasing concentrations of MSI-2 intensified the observed band shift to the larger RNA/Protein complex and decreased the equivalent unbound RNA band. The interaction between a 1:4 ratio of *RNA^WT^:MSI-2 was competed with increasing concentrations of unlabelled **B)** RNA^WT^ or **C)** RNA^mut^ (_63_CAACUU_68_-mut). **D)** Band shifts in the unlabeled RNA competition EMSAs were quantified by densitometry and expressed as % change in the density of the RNA/Protein complex bands, normalized to the equivalent total lane density, for each competition ratio and compared each time to ratio 1:0. N=3, error bars represent standard error from the mean and significance was measured by two-tailed T-test (* = P < 0.05, ** = P < 0.01, *** = P < 0.001). Grey block triangles indicate increasing concentrations of specific reactants.

To further investigate the specificity and location of the interaction, the putative MSI-2 binding site _63_AUUAAU_68_ was mutated to _63_CAACUU_68_ (henceforth termed _63_CAACUU_68_-mut), which was predicted *by in silico* analysis not to affect local RNA structure (Supplementary data 4A and B). EMSA competition, with unlabelled _63_CAACUU_68_-mut RNA-330, was less efficient at competing for MSI-2 8-193 binding than wild-type CHIKV RNA (Fig 2C and D). These EMSA results are consistent with an interaction between MSI-2 and the upstream 1-330 nt region of the CHIKV genome. Furthermore, reduced competition with mutant _63_CAACUU_68_-mut compared with wild type, was consistent with a specific interaction, in which nts _63_AUUAAU_68_ function as an MSI binding motif.

### Replication of CHIKV infectious virus and sub-genomic replicon was restricted by MSI small molecule inhibitor Ro 08-2750

In order to investigate the potential role of MSI on CHIKV replication, we first assessed the effect of a well-characterised MSI small molecule inhibitor, Ro 08-2750, on CHIKV productive replication in RD cells using both CHIKV infectious virus and a sub-genomic replicon (SGR) assay (Fig 3A). A number of studies have demonstrated that Ro interacts with the MSI RNA binding domains and acts as a competitive inhibitor for its RNA binding activity and subsequent MSI functions within the cell (22,23) (24). In the current study Ro 08-2750 was used at a maximum non-toxic dose as determined by MTT assay (Supplementary data 5).

**Figure 3.**
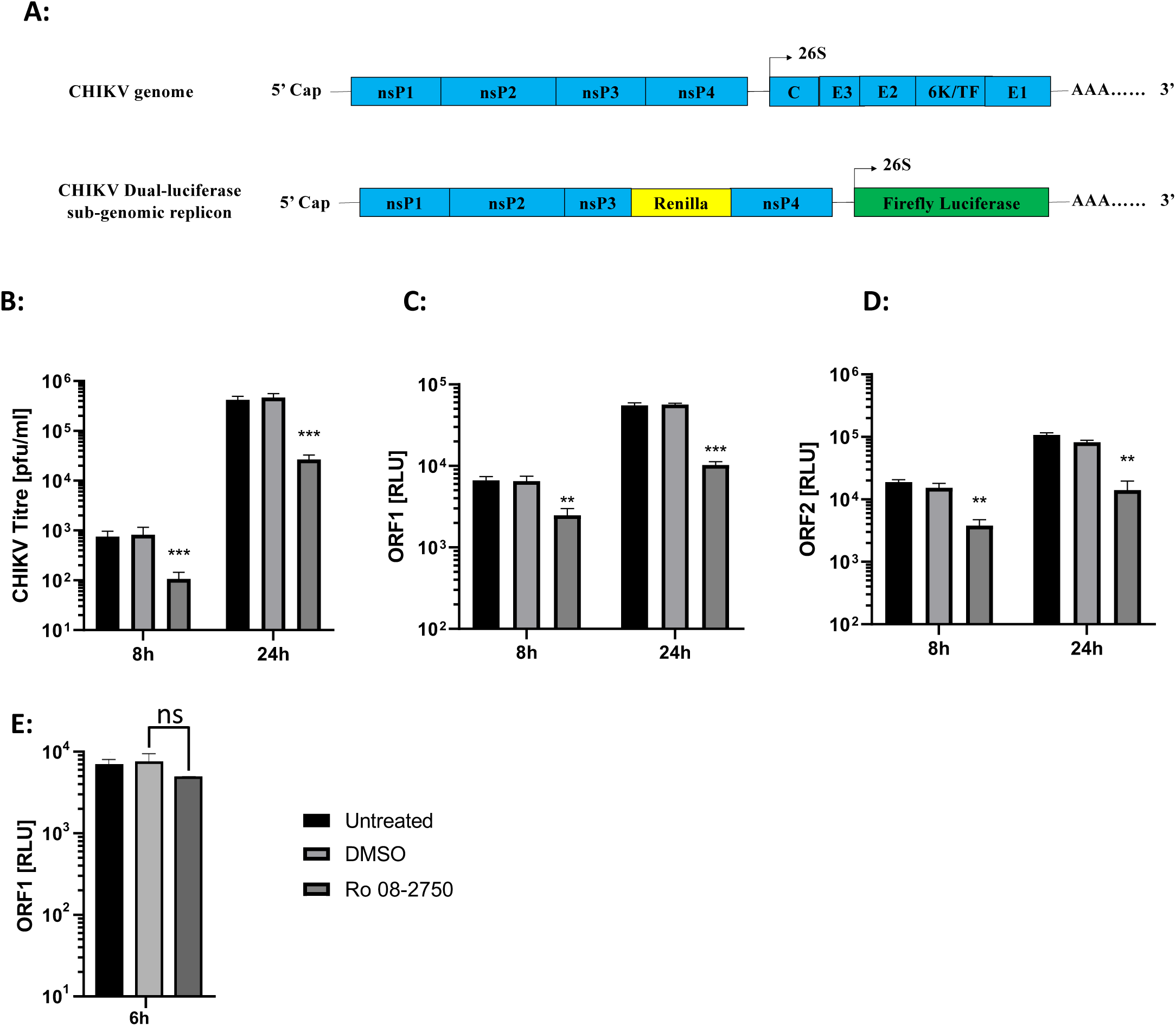
Ro 08-2750 signifyingly inhibits replication of infectious CHIKV and the CHIKV-SGR **A)** Schematic representations of CHIKV infectious clone (top) compared to the sub-genomic replicon (SGR) (bottom) in which a *Renilla* luciferase (RLuc) reporter gene is fused within the nsp3 coding sequence and the structural genes of ORF-2 are replaced by a firefly luciferase (Fluc) reporter gene. Replication is expressed in Relative Light Units [RLU] **B)** Ro significantly inhibits productive CHIKV productive replication relative to DMSO treated negative controls at 8 and 24 hpi. **C)** and **D**) Ro 08-2750 significantly inhibits CHIKV-SGR replication, measured by both ORF-1 and ORF-2 expression, relative to DMSO treated negative controls at 8 and 24 hpt. **E)** Ro 08-2 750 had no significant (ns) effect on CHIKV-SGR (GDD-GAA) translation, measured by ORF-1 expression. N=3, error bars represent standard error from the mean and significance was measured by two-tailed T-test (* = P < 0.05, ** = P < 0.01, *** = P < 0.001).

To assay the effect of Ro 08-2750 on early and late stages of CHIKV replication, productive CHIKV infection was measured at 8 and 24 hpi. Pre-treatment and incubation in the presence Ro significantly inhibited infectious CHIKV replication by ∼10-fold, relative to DMSO treated negative controls (Fig 3B). In order to investigate the effect of Ro 08-2750 on specific stages of CHIKV replication we used an SGR construct, in which ORF-2 was replaced by a firefly luciferase (FLuc) gene and a *Renilla* luciferase (Rluc) gene was fused in-frame with ORF-1 nsP3 (Fig 3A)(10). The SGR assay enabled us to measure the effect of Ro 08-2750 on CHIKV genome replication and translation, in isolation of other stages of virus infection, such as entry or egress. SGR ORF-1 and ORF-2 expression was significantly inhibited at both 8 and 24 hpt, relative to DMSO treated negative controls, indicating that Ro 08-2750 inhibited CHIKV replication at the level of virus genome replication or translation (Fig 3C).

In order to investigate the possibility that Ro 08-2750 specifically inhibited ORF-1 translation, we used a replication incompetent SGR, incorporating an nsP4 GDD>GAA substitution in the RdRp active site (CHIKV-SGR (GDD>GAA)). Such an approach enabled the efficiency of ORF-1 translation to be measured by *Renilla* luciferase expression, in the presence and absence of Ro 08-2750, under conditions in which genome replication could not occur. Translation of input SGR 5ʹcapped transcripts was measured at 6 hpt in RD cells (Fig 3E). No significant difference in level of translation was observed between the CHIKV-SGR (GDD>GAA) in the presence or absence of Ro 08-2750. These results indicate that Ro 08-2750 did not influence ORF-1 translation but inhibited SGR genome replication.

### Ro inhibited CHIKV *trans*-complementation assay

Inhibition of SGR replication demonstrated that Ro was inhibiting CHIKV replication at the level of virus genome replication or translation. In order to dissect this further, we utilised a *trans*-complementation assay, that enabled the measurement of CHIKV genome replication in isolation of virus translation (Fig 4A)(25). The *trans*-complementation system utilised a CHIKV replicase-expressing plasmid (pCHIKV-nsP1234). pCHIKV-nsP1234, expresses the virus nsPs from a CMV promoter and is codon optimised to disrupt RNA sequence motifs, such as protein binding signals, and the formation of wild-type CHIKV RNA structure. The expressed nsPs replicate a CHIKV reporter construct (pCHIKV-FLuc/GLuc), in which the majority of ORF-1 is replaced by an FLuc gene and ORF-2 by the Gaussia luciferase (GLuc) gene. The effect of Ro on pCHIKV-FLuc/GLuc expression was measured at 8 and 24 hpt in cells pre-treated with and maintained in the presence of Ro. Both ORF-1 and ORF-2 expression was observed to be significantly inhibited relative to the DMSO treated negative controls, indicating that Ro was specifically inhibiting CHIKV genome replication.

**Figure 4.**
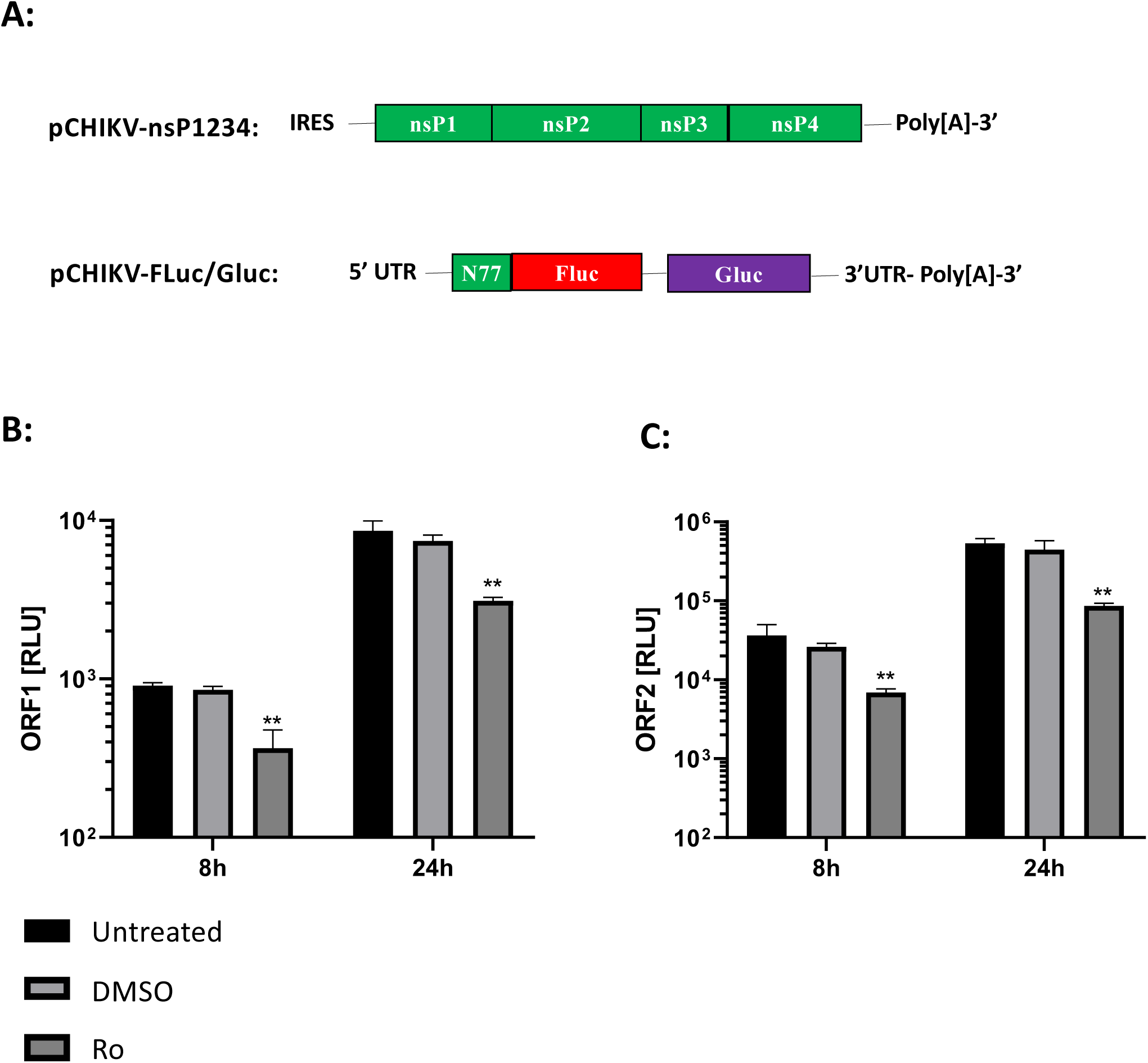
Ro signifyingly inhibits CHIKV genome replication. **A)** Schematic representation of CHIKV *trans-*complementation assay showing codon optimised pCHIKV-nsP1234 (top) from which the CHIKV nsPs were translated and pCHIK-Fluc/Gluc (bottom) in which ORF-1 was replaced by an Fluc reporter gene, fused to the first 77 nts or CHIKV ORF-1 (N77) downstream of the authentic CHIKV 5’UTR. ORF-2, flanked by the authentic intragenic (SG) and 3’ UTRs, was replaced by a Gluc reporter gene. **B)** and **C)** Ro significantly inhibited CHIKV genome replication of the *trans-*complementation assay, measured by both ORF-1 and ORF-2 expression, relative to DMSO treated negative controls at 8 and 24 hpt. N=3, error bars represent standard error from the mean and significance was measured by two-tailed T-test (* = P < 0.05, ** = P < 0.01, *** = P < 0.001).

### MSI-2 shRNA silencing inhibits infectious CHIKV replication

In order to confirm that Ro induced inhibition of CHIKV genome replication was due to specific inhibition of MSI, rather than an unrecognised off target effect of Ro, we next investigated CHIKV replication following shRNA silencing of MSI expression. Analysis of RD cell total protein extract by western blot, demonstrated that MSI-2 was strongly expressed in RD cells while MSI-1 was expressed at a very low level (Supplementary data 6). Consequently, we initially investigated the effect of specific MSI-2 shRNA silencing on CHIKV replication. RD cells were transduced with lentiviral vectors encoding shRNA against human MSI-2 and successful silencing was confirmed by western blot (Fig 5A). Negative control cells were treated with either transfection reagent only (mock) or non-specific scrambled shRNA. Following infection of MSI-2 depleted and negative control cells, productive CHIKV infection was measured by plaque assay at 8 and 24 hpi (Fig 5B). Strand specific qRT-PCR was also used to measure levels of genomic (positive strand) and replication intermediate (negative strand) CHIKV RNA at both time points (Fig 5C and D). As observed previously following treatment with Ro, relative to the scrambled shRNA control productive CHIKV replication was significantly inhibited following shRNA MSI-2 silencing (Fig 5B). Similarly, strand specific qRT-PCR demonstrated that both genomic and replication intermediate CHIKV RNA levels were significantly inhibited (Fig 5C and D). These results are consistent with CHIKV requiring MSI-2 for efficient genome replication.

**Figure 5.**
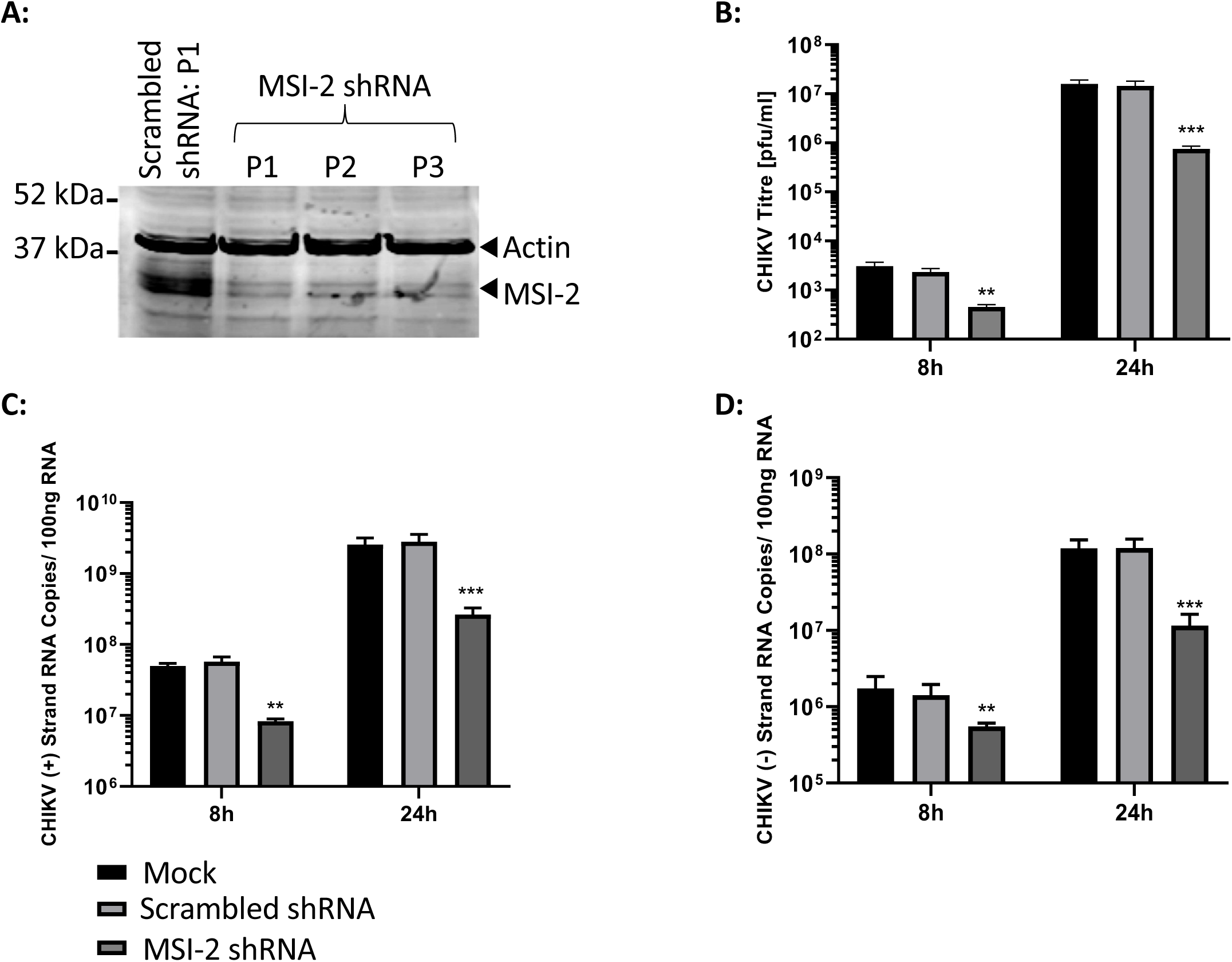
shRNA suppression of MSI-2 significantly inhibits replication of infectious CHIKV. **A)** Western blot analysis of total cellular protein extracted from Rd cells and compared to negative control scrambled shRNA, demonstrated consistent shRNA knockdown of MSI-2 over 3 serial passages (P1-P3). MSI-2 suppression significantly inhibited productive CHIKV replication, relative to scrambled shRNA at 8 and 24 hpi measured by plaque assay **(B)** and strand specific qRT-PCR for the the virus genomic **(C)** and negative intermediate **(D)** strands. N=3, error bars represent standard error from the mean and significance was measured by two-tailed T-test (* = P < 0.05, ** = P < 0.01, *** = P < 0.001).

### MSI-2 is required for efficient CHIKV genome replication

In order to confirm that MSI-2 silencing had the same effect on CHIKV genome replication as Ro small molecule inhibition, we repeated analysis with the SGR and *trans*-complementation systems, following shRNA silencing of MSI-2. As previously described, MSI-2 shRNA silenced RD cells were transfected with the SGR system and ORF-1 and ORF-2 expression measured by RLuc and FLuc expression at 8 and 24 hpt (Fig 6A and B). Expression of both ORF-1 and ORF-2 was significantly inhibited at both 8 and 24 hpt, consistent with an MSI-2 requirement for CHIKV genome replication or translation. Analysis at the same time points using the *trans*-complementation system, following shRNA MSI-2 silencing and measuring ORF1 and ORF2 expression, confirmed significant inhibition of CHIKV genome replication when measured in isolation of the effects of other stages of the virus replication cycle (Fig 6C and D)

**Figure 6.**
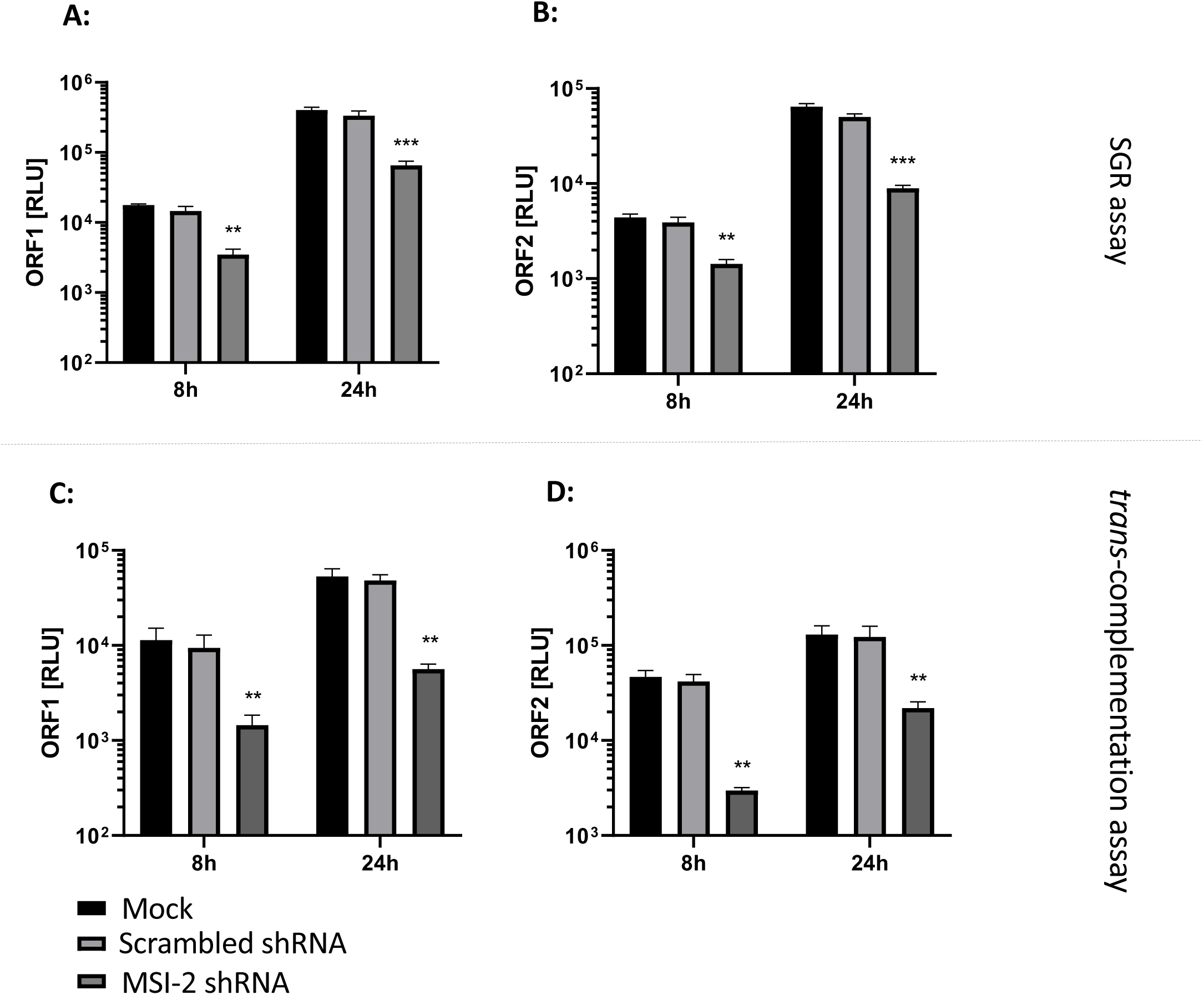
shRNA suppression of MSI-2 significantly inhibits CHIKV-SGR replication and CHIKV genome replication. **A)** and **B)** shRNA suppression of MSI-2 significantly inhibited CHIKV-SGR replication, measured by both ORF-1 and ORF-2 expression, relative to scrambled shRNA negative controls at 8 and 24 hpt. **C)** and **D)** shRNA suppression of MSI-2 significantly inhibited CHIKV genome replication of the *trans-* complementation assay, measured by both ORF-1 and ORF-2 expression, relative to DMSO treated negative controls at 8 and 24 hpt. N=3, error bars represent standard error from the mean and significance was measured by two-tailed T-test (* = P < 0.05, ** = P < 0.01, *** = P < 0.001).

### Both MSI-2 and MSI-1 have a proviral effect on CHIKV genome replication

While results clearly demonstrated that MSI-2 was required for efficient CHIKV replication, it remained unclear if MSI homologue MSI-1 was also agonistic for CHIKV replication. As previously described, western blot analysis showed that MSI-1 was not highly expressed in RD cells. Consequently, we analysed MSI homologue redundancy in Huh7 human hepatoma cells. MSI-1 and MSI-2 homologues are strongly expressed in Huh7 cells (Supplementary data 6) and they are highly permissive for CHIKV replication (26). Following individual and combined siRNA silencing of MSI-1 and MSI-2 in Huh7 cells (Supplementary data 7), CHIKV replication was assayed by plaque assay and ssRT-qPCR at 8 and 24 hpi (Fig 7A, B and C). As was observed for RD cells, MSI-2 silencing significantly inhibited CHIKV replication, measured by both plaque assay and strand specific ssRT-qPCR. Interestingly, it was observed that MSI-1 silencing in Huh7 cells had a similar significant inhibitory effect and that siRNA co-silencing of both MSI-1 and MSI-2 had a synergistic effect on inhibition of CHIKV replication. In concordance with previous results, siRNA silencing of MSI-2 in RD cells significantly inhibited CHIKV replication, by comparable levels observed following shRNA silencing and inhibition by Ro. Consistent with low MSI-1 expression in RD cells, siRNA co-silencing of both MSI-1 and MSI-2 did not significantly increase levels of CHIKV inhibition (Fig 7D, E and F).

**Figure 7:**
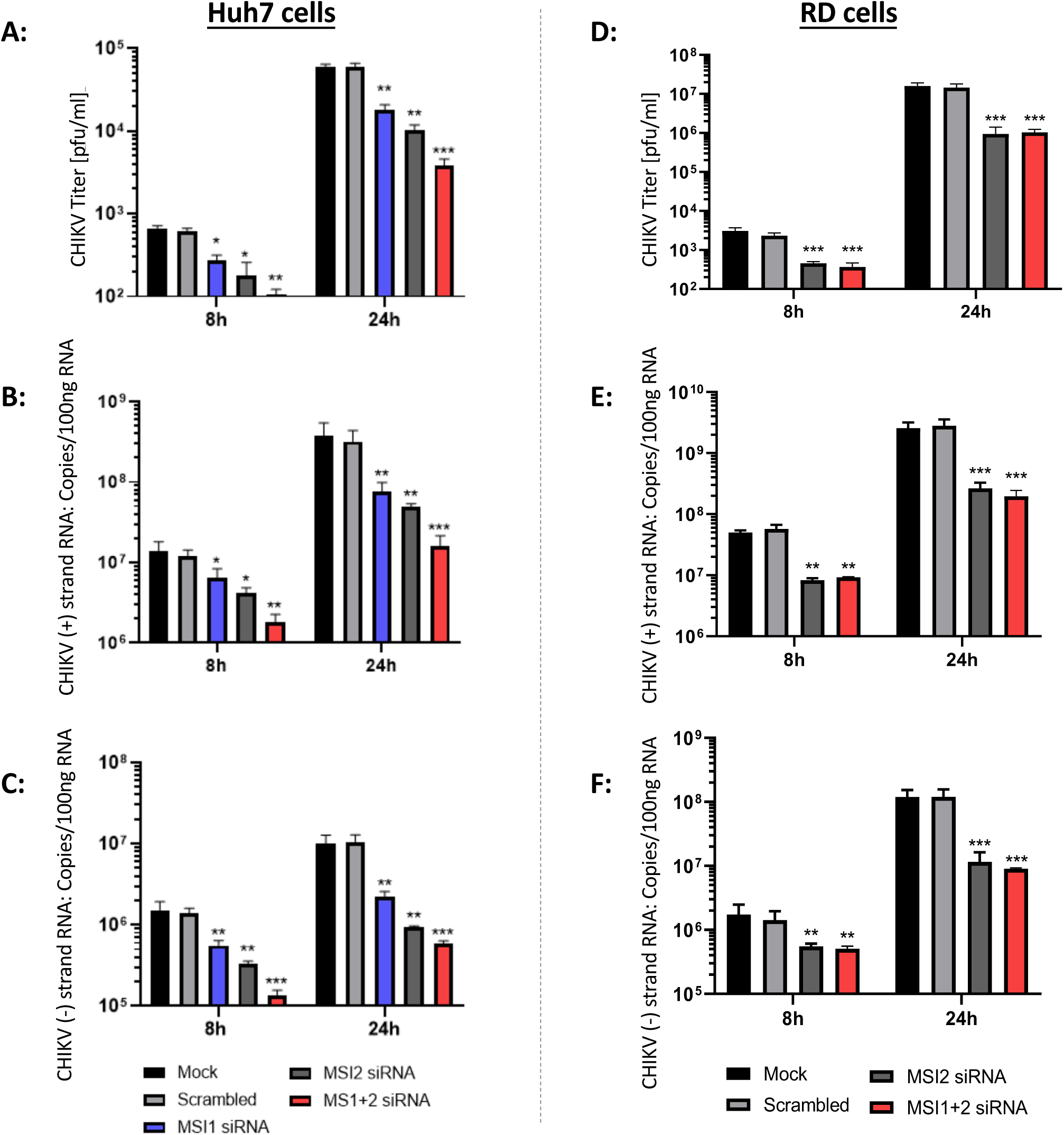
siRNA Depletion of either MSI-1 or MSI-2 significantly inhibited CHIKV replication in Huh7 cells and co-depletion of both MSI-1 and MSI-2 had a synergistic effect on CHIKV inhibition in Huh7 cells. siRNA depletion of MSI-2 significantly inhibited CHIKV replication in RD cells and co-depletion of both MSI-1 and MSI-2 did not increase the level of CHIKV inhibition. siRNA depletion of MSI-1 and MSI-2 significantly inhibited CHIKV replication in Huh7 cells relative to scrambled siRNA at 8 and 24 hpi measured by plaque assay **(A)** and strand specific qRT-PCR for the the virus genomic **(B)** and negative intermediate **(C)** strands. siRNA depletion of MSI-2 in RD cells significantly inhibited CHIKV replication in relative to scrambled siRNA at 8 and 24 hpi measured by plaque assay **(D)** and strand specific qRT-PCR for the the virus genomic **(E)** and negative intermediate **(F)** strands. N=3, error bars represent standard error from the mean and significance was measured by two-tailed T-test (* = P < 0.05, ** = P < 0.01, *** = P < 0.001).

### Reverse genetic analysis of _63_AUUAAU_68_ MSI binding site

In order to further investigate the role of the _63_AUUAAU_68_ MSI binding motif in CHIKV replication, we took a reverse genetic approach, in which the _63_AUUAAU_68_>_63_CAACUU_68_ mutations were incorporated into both the *trans*-complementation assay and infectious CHIKV. While _63_AUUAAU_68_>_63_CAACUU_68_ disrupted the MSI binding site, it was predicted to have no effect on local RNA structure (Supplementary data 4A and B). Similar to previous results following MSI inhibition, expression of both ORF-1 and ORF-2 from the *trans*-complementation system were significantly inhibited in _63_CAACUU_68_-mut relative to the wild type at 8 and 24 hpt (Fig 8A and B); indicating that mutation of the putative MSI binding site significantly inhibited CHIKV at the level of virus genome replication. Interestingly, despite repeated attempts, we were not able to rescue infectious CHIKV _63_CAACUU_68_-mut virus, indicating that disruption of the potential MSI binding site completely inhibited infectious CHIKV replication (Fig 8C). In further reverse genetic analysis, we analysed the effect of substitution _A_67_G_ on infectious CHIKV replication. _A_67_G_ modified the _63_AUUAAU_68_ to match the published MSI canonical binding site but disrupted adjacent downstream RNA structure (Supplementary data 4C). Following infection at MOI 0.1 in RD cells, CHIKV _A_67_G_ mutant and wild-type control virus was harvested at 8 and 24 hpi and titred by plaque assay (Fig 8D). Compared to wild-type, mutant _A_67_G_ significantly inhibited CHIKV replication at 8 and 24 hpi, indicating that nucleotides _63_AUUAAU_68_ are constrained by both a requirement to interact with MSI-2 and also to maintain essential local RNA structure.

**Figure 8.**
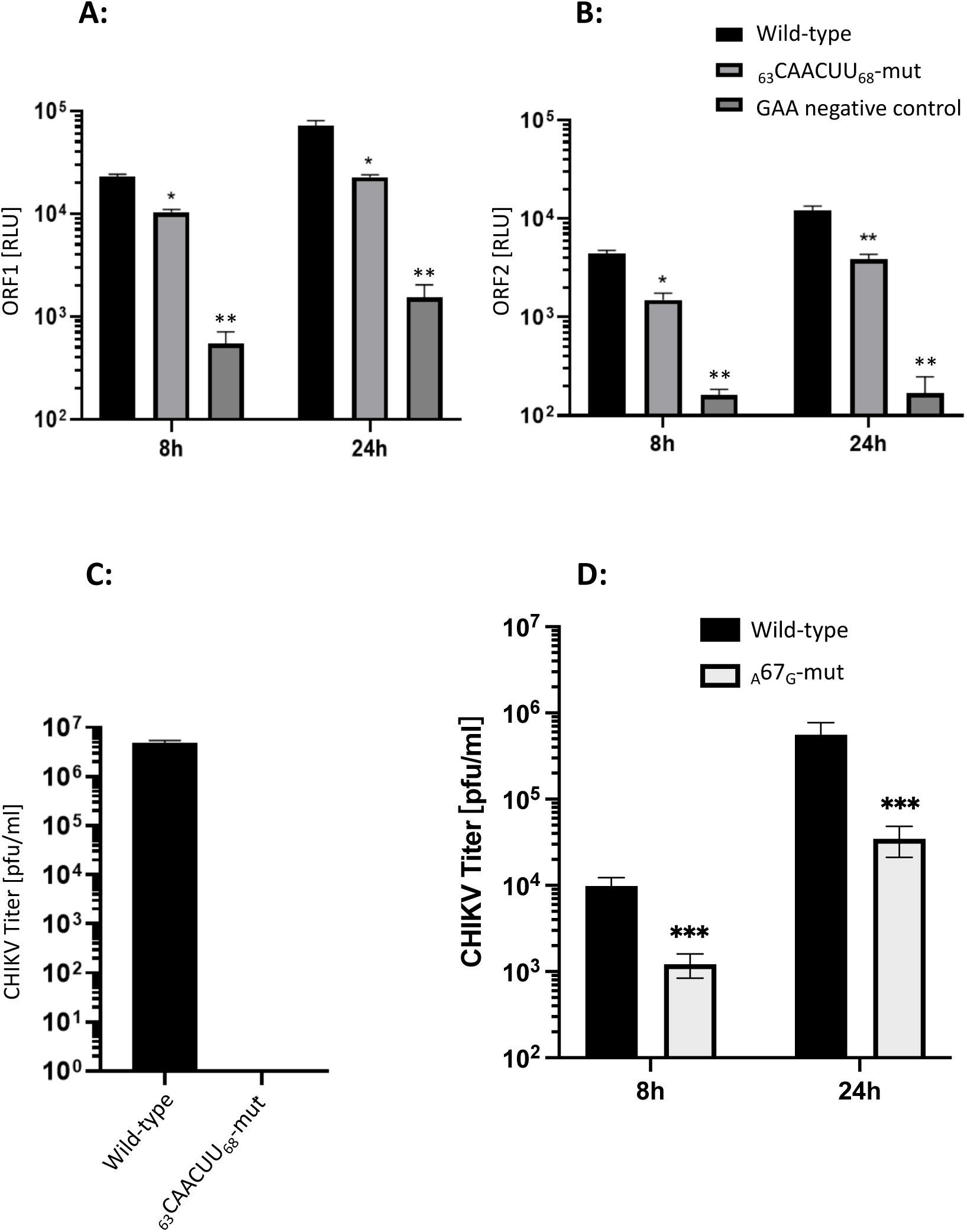
Substitutions disrupting the predicted MSI binding site (_63_CAACUU_68_-mut) but maintaining local RNA structure prevented virus rescue and significantly inhibited CHIKV genome replication. Substitution _A_67_G_ maintained the predicted MSI-2 binding motif but disrupted local RNA structure and significantly inhibited CHIKV replication. **A)** and **B)** Mutation _63_CAACUU_68_ significantly inhibited CHIKV genome replication of the *trans-*complementation assay, measured by both ORF-1 and ORF-2 expression, relative to wild-type positive controls at 8 and 24 hpt. **C)** Mutation _63_CAACUU_68_-mut prevented rescue of CHIKV following transfection of capped *in vitro* transcribed RNA into BHK cells. Released virus was measured by plaque assay of supernatant 24 hpt and compared to positive control wild-type infectious CHIKV *in vitro* transcribed RNA, which was transfected and analysed in parallel. **D)** Mutation ^A^67^G^ significantly inhibited CHIKV replication at 8 hpi and 24 hpi relative to wild type CHIKV following infection of RD cells (MOI 0.1). N=3, error bars represent standard error from the mean and significance was measured by two-tailed T-test (* = P < 0.05, ** = P < 0.01, *** = P < 0.001).

### Ro 08-2750 further inhibited CHIKV genome replication in _63_CAACUU_68_-mut

In previous experiments, we demonstrated that mutations to disrupt the _63_AUUAAU_68_ 5’UTR MSI binding site or MSI-2 depletion/inhibition restricted CHIKV genome replication. In order to investigate the potential synergistic effects of these approaches we repeated analysis of _63_CAACUU_68_-mut *trans*-complementation assay replication, in the presence of Ro 08-2750 (Fig 9). Interestingly, in the presence of Ro 08-2750, _63_CAACUU_68_-mut replication was inhibited to a significantly greater level than _63_CAACUU_68_-mut in the absence of Ro 08-2750, at both 8 and 24 hpt. These results are consistent with MSI-2 acting through interaction with _63_AUUAAU_68_ in the 5’UTR and a further region of the CHIKV genome present in the sub-genomic *trans* complementation reporter (i.e., 5’UTR, intragenic region or 3’UTR).

**Figure 9.**
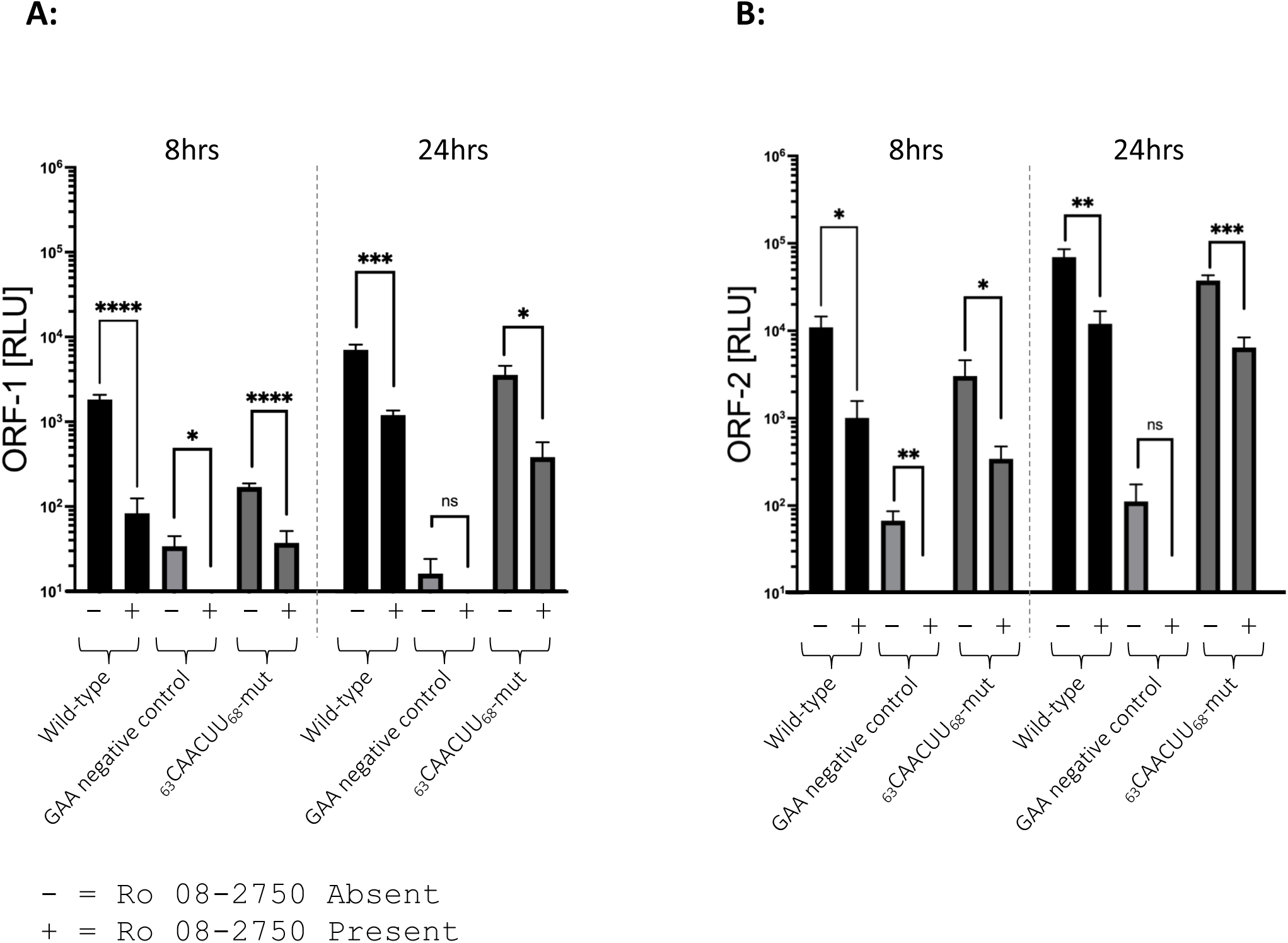
Combining substitutions within the predicted MSI binding site (_63_CAACUU_68_-mut) and Ro 08-2750 had a synergistic affect, significantly increasing inhibition of CHIKV genome replication. **A)** and **B)** Mutation of the MSI binding site combined with Ro 08-2750 significantly increased inhibition of the CHIKV *trans-* complementation assay, measured by both ORF-1 and ORF-2 expression, relative to _63_CAACUU_68_-mut in the absence of Ro 08-2750 at 8 and 24 hpt. N=3, error bars represent standard error from the mean and significance was measured by two-tailed T-test (* = P < 0.05, ** = P < 0.01, *** = P < 0.001).

## DISCUSSION

For the first time, in this study we demonstrate that cellular RNA binding protein MSI is required for efficient CHIKV replication. Our data demonstrates that MSI-2 interacts directly with the 5’ end of the CHIKV genome and is consistent with an interaction at position _63_AUUAAU_68_. A single-stranded region located between two conserved stem-loops and immediately upstream of the AUG start codon (Fig 1) (10). Inhibition of MSI-2 expression by siRNA or shRNA, its RNA-binding activity by Ro 08-2750 and reverse genetic disruption of the CHIKV-5’UTR/MSI-2 interaction, inhibited CHIKV replication at the level of virus genome replication. In further analysis, we demonstrated that depletion of both MSI-1 and MSI-2 homologues resulted in a synergistic increase in the level of CHIKV inhibition, supporting the premise that both MSI homologues have a redundant pro-viral effect on CHIKV replication.

While depletion of MSI-2 by siRNA and shRNA clearly demonstrated significant inhibition of CHIKV replication, measured by infectious virus production, CHIKV-SGR replication or a *trans*-complementation assay, we did not observe complete inhibition of virus replication. Western blot analysis indicated that we did not achieve complete ablation of MSI-2 expression by either siRNA or shRNA, presumably reducing the level of CHIKV inhibition observed. We demonstrated by western blot that in RD cells MSI-2 is expressed to a high level but the MSI-1 homologue is only expressed to a low level. However, given the synergistic effect that we observed when silencing expression of both MSI-2 and MSI-1 in Huh7 cells (in which both are expressed to high levels), the ability of MSI homologues to complement for each other may also have reduced the level of CHIKV inhibition observed. This hypothesis is consistent with reverse genetic results, in which _63_AUUAAU_68_> **_6_**_3_CAACUU_68_ mutation of the MSI binding site, dramatically inhibited CHIKV-5’UTR/MSI-2 binding affinity and completely prevented rescue of infectious mutant virus.

Mutation _63_AUUAAU_68_> **_6_**_3_CAACUU_68_ also significantly inhibited replication in the *trans-*complementation assay. However, genome replication was not completely ablated. These differences may indicate that this region has a further function during infectious virus replication, that is not assayed in the *trans-*complementation assay, such as during virion assembly. Further reverse genetic analysis, in which substitution _A_67_G_ engineered a canonical MSI binding site but was predicted to disrupted downstream RNA structure, significantly inhibited infectious CHIKV replication, suggesting that _63_AUUAAU_68_ is constrained by both a requirement to bind MSI and to maintain local RNA structure.

Translation and replication of positive-sense RNA virus genomes, such as those of CHIKV are mutually exclusive processes – with translation initiating at the 5’ end of the genome and replication at the 3’. Consequently, it is essential that such viruses have mechanisms for temporal control of both processes. In many such viruses (e.g. Hepatitis C virus and Polio virus) control involves dynamic interactions between RNA structures in the virus genome and host/virally expressed protein complexes (8). The mechanisms and interactions which CHIKV and other alphaviruses use for temporal control of genome translation and replication remain unclear. However, a recent study suggested a crucial role for the cellular helicase DHX9. Matkovic et al, demonstrated that interaction between DHX9 and the 5’ end of the virus genome upregulates CHIKV ORF-1 non-structural protein translation, while simultaneously inhibiting replication of its genome (27). Build-up of nsP2 caused proteosome induced degradation of DHX9, although the detailed mechanism for this remains unclear.

While, genome replication of positive-sense RNA viruses initiates at the 3’ end, it is commonly controlled by promoter elements and interactions with RNA binding proteins at the 5’ end. Results described in this and previous studies are consistent with a model in which temporal control of CHIKV ORF-1 translation and genome replication is controlled by a mechanism involving interactions between DHX9, MSI-2 and the 5’ region of the virus genome. However, the dynamics and mechanism by which the opposing roles if MSI-2 and DHX-9 influence CHIKV replication and their interactions with other regions of the virus genome remains unclear. For example, EMSA and reverse genetic results presented here clearly demonstrate that MSI-2 binds to nucleotides _63_AUUAAU_68_ in the 5’UTR and that this interaction is required for efficient CHIKV genome replication. However, the synergistic effect of Ro 08-2750 and _63_CAACUU_68_-mut are consistent with MSI-2 also interacting with another region of the CHIKV genome. Bioinformatic analysis has not predicted further MSI binding motifs within the CHIKV genome. Consequently, we hypothesise that any such alternative interactions may be indirect, via a protein complex and are the focus of ongoing studies.

In summary, using MSI-2 depletion and/or Ro 08-2750 small molecule inhibitor we demonstrate that MSI-2 RNA binding protein is a critical host factor for efficient CHIKV replication but is not required for ORF-1 translation. Inhibition of ORF-1 and ORF-2 signal from the *trans*-complementation assay indicate that MSI-2 is required at the level of CHIKV genome replication. Furthermore, EMSA and reverse genetic results are consistent with a direct interaction between MSI-2 and _63_AUUAAU_68_ in the positive genomic strand of the virus 5’UTR, suggesting a role in negative strand synthesis. Results from this study are important both for our understanding of the fundamental interactions essential to replication of this important human pathogen and for future studies towards specific antiviral therapies.

## Data availability statement

The data underlying this article will be shared on reasonable request to the corresponding author.

---

**Supplementary data 1:**
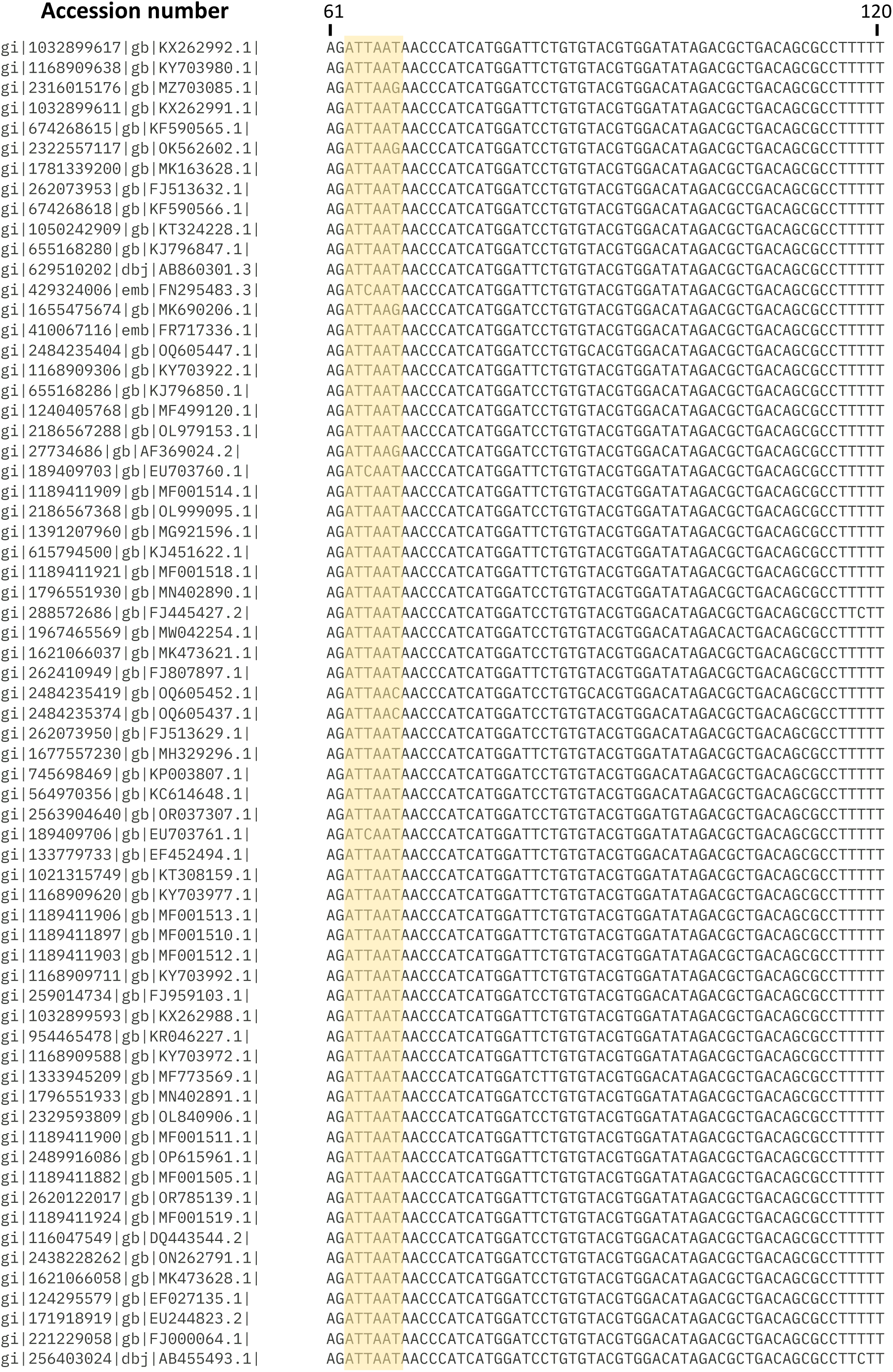
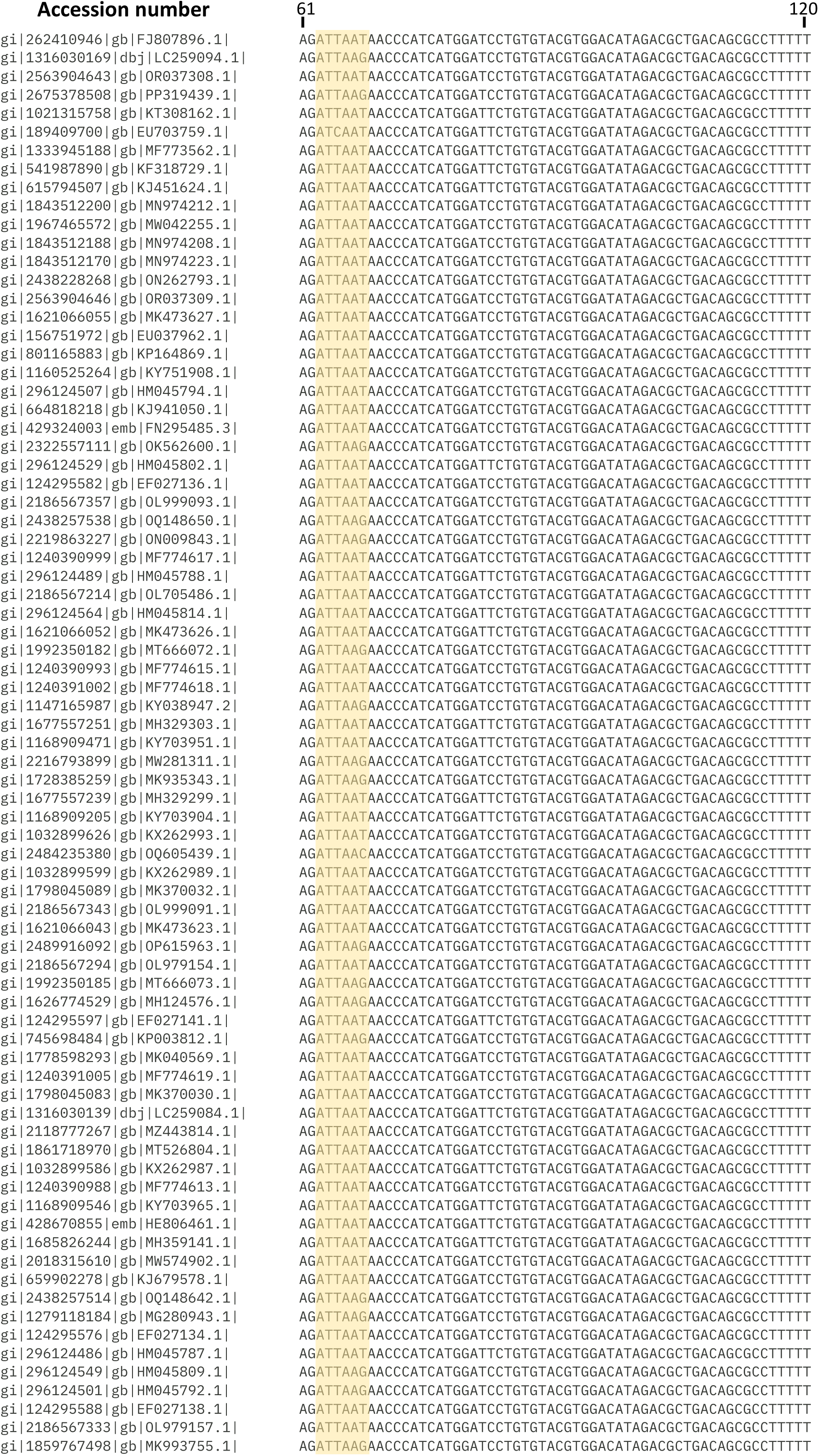
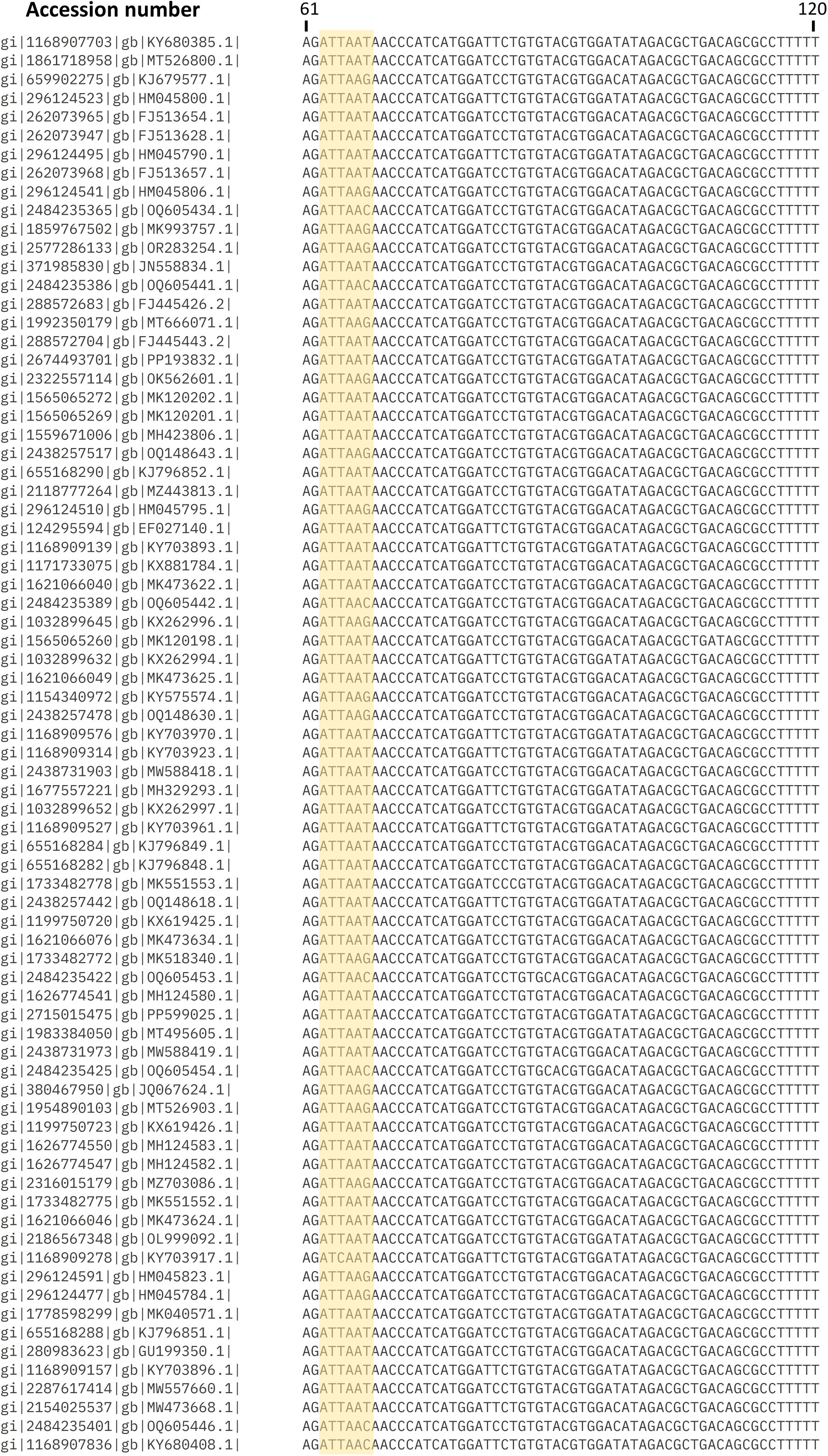
Alignment of CHIKV 5’ region (nts 61-120) showing variation within 5’UTR MSI-2 binding site. All complete CHIKV genomes were downloaded from the NCBI, by using the selection for complete genomes and search term for CHIKV. Duplicates were removed, leaving one representative of each, which were aligned using CLUSTAL OMEGA (multiple sequence alignment) on EBI default settings. MSI-2 binding site indicated by orange box.

**Supplementary data 2:**
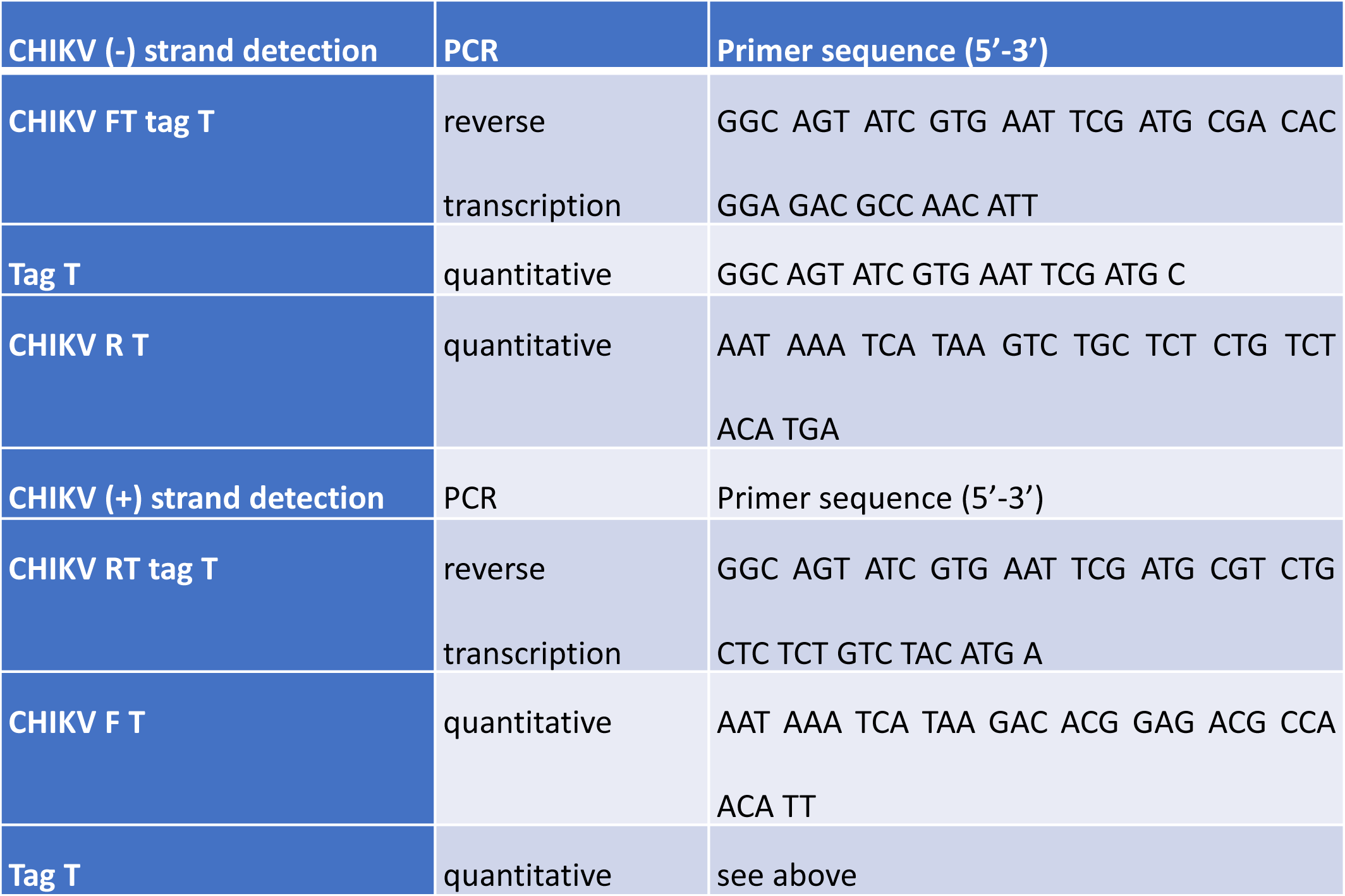
Primers for the reverse transcription and quantitative PCRs for CHIKV strand-specific detection.

**Supplementary data 3:**
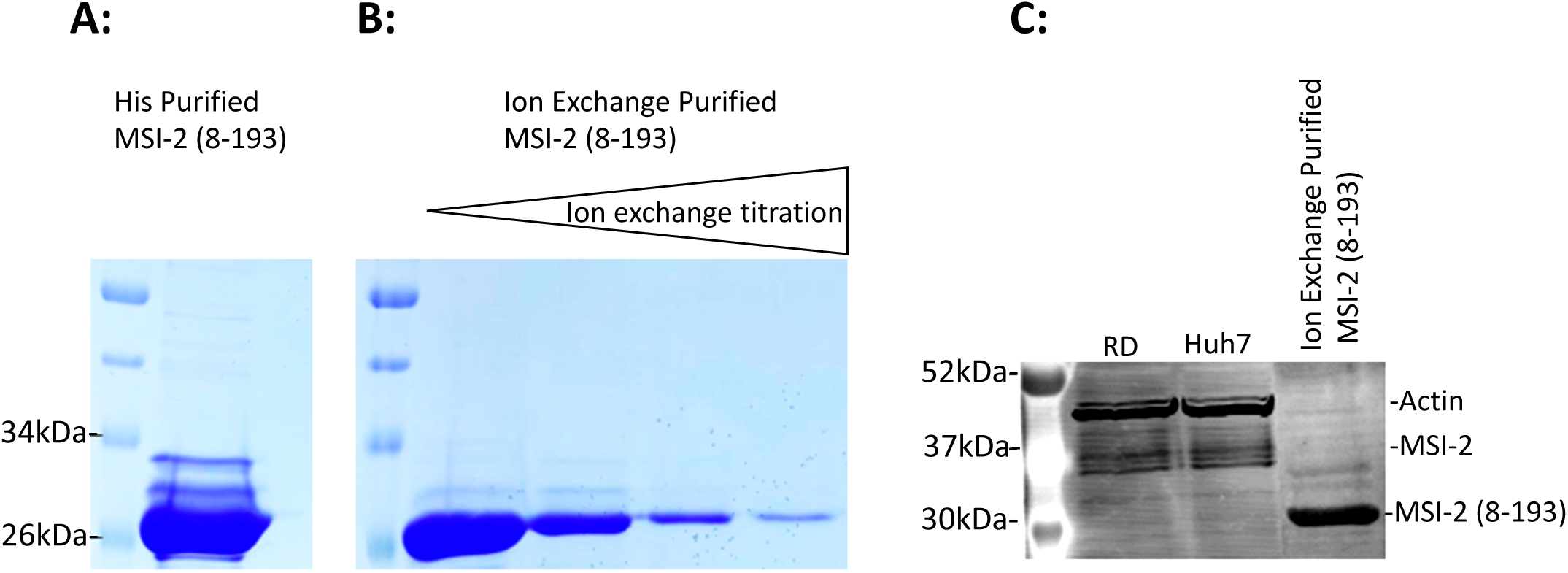
Expression and purification of MSI-2 Coomassie stained PAGE analysis following **A)** His Tag and **B)** Ion exchange chromatography. **C)** Ion exchange purified MSI-2 (8-193) analyzed by western blot, relative to total protein extracted from RD and Huh7 cells.

**Supplementary data 4:**
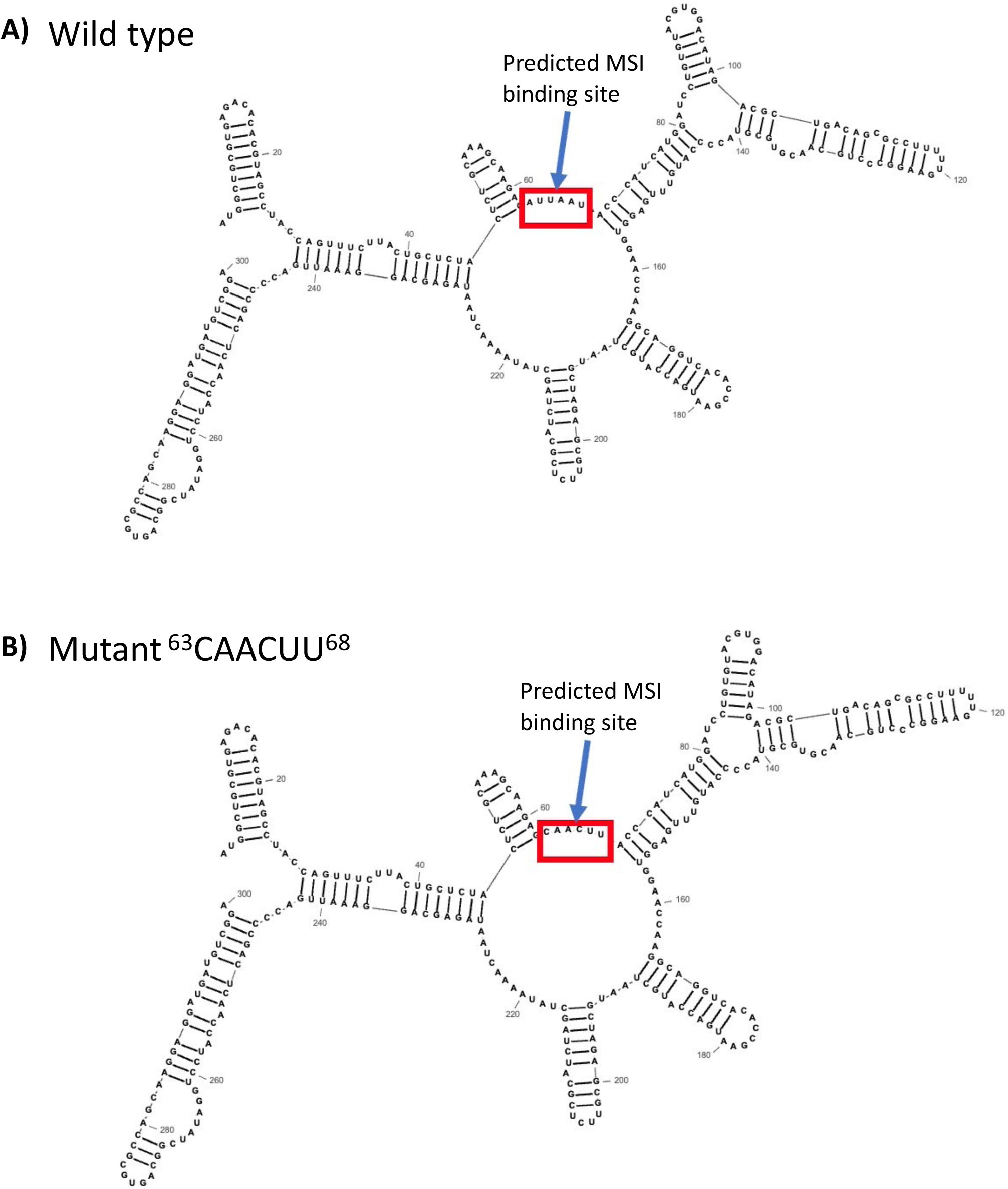
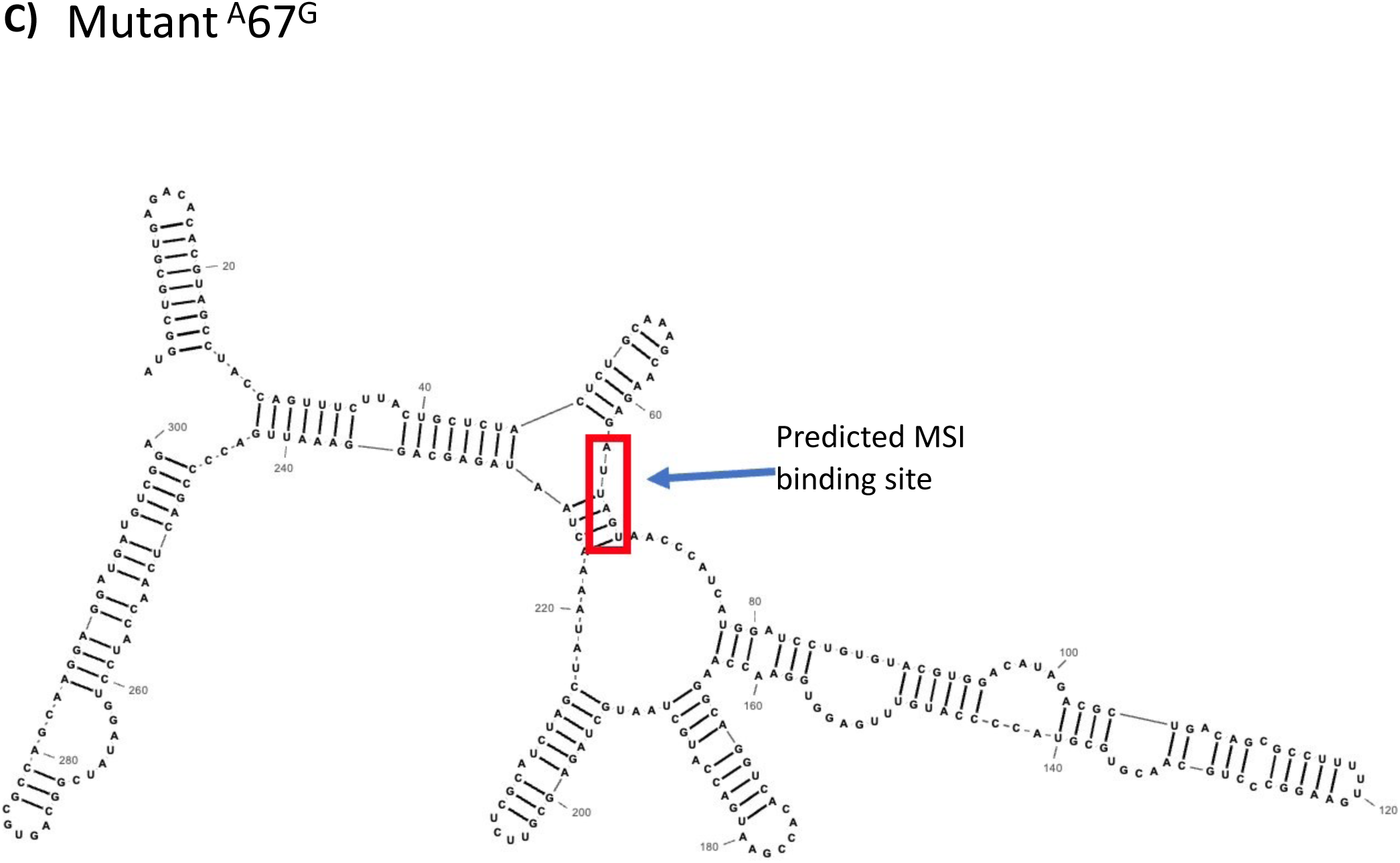
CHIKV predicted RNA structure (nts 1-300) in **A)** wild type **B) m**utant ^63^CAACUU^68^ and **C)** mutant ^A^67^G^. RNA structure mapped free energy minimization using Pfold on default settings.

**Supplementary data 5:**
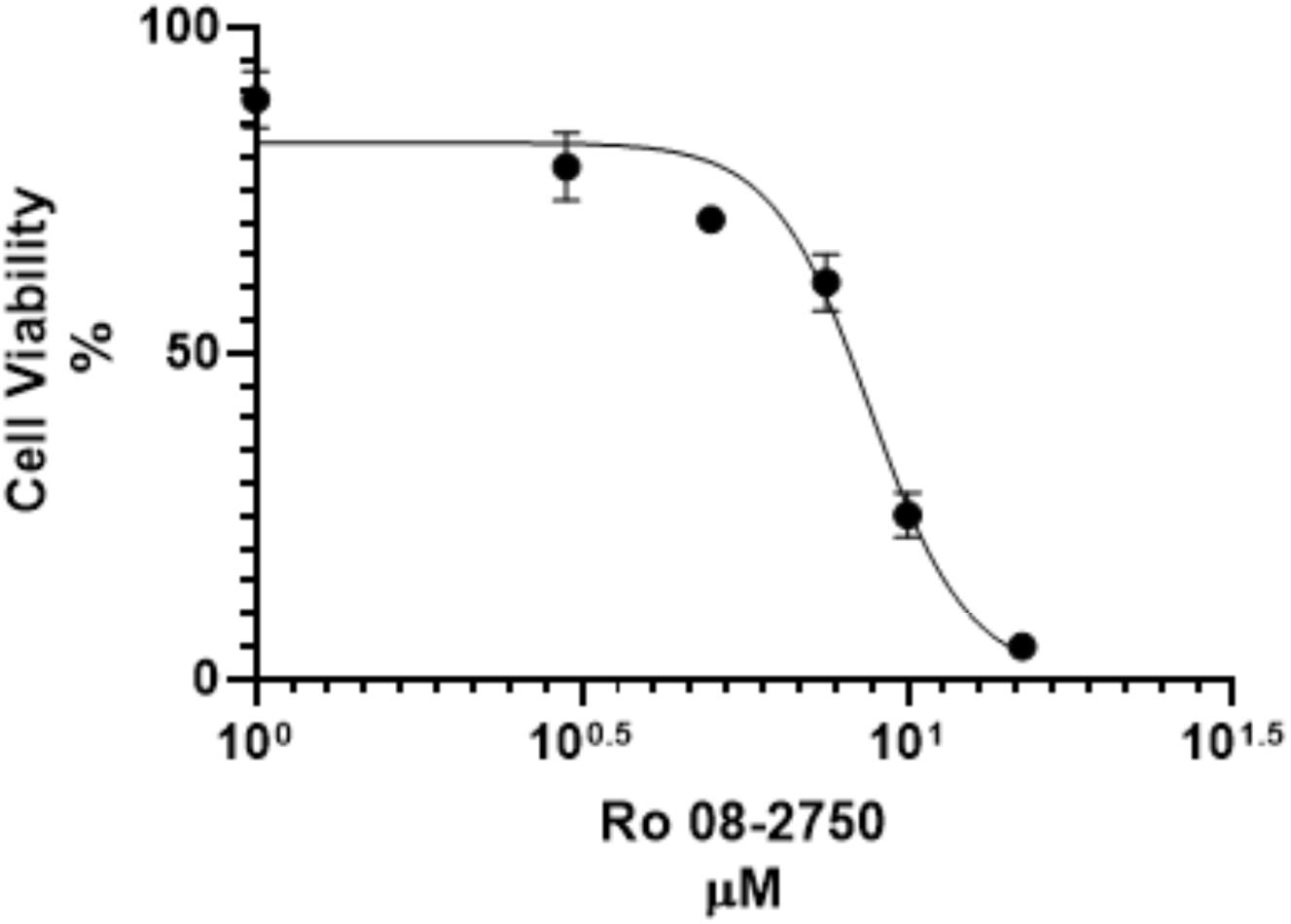
MTT cytotoxicity assay for Ro 08-2750 in RD cells across a titration of 0, 05, 1, 3, 5 10 and 20uM. N=3, error bars represent standard error from the mean.

**Supplementary data 6:**
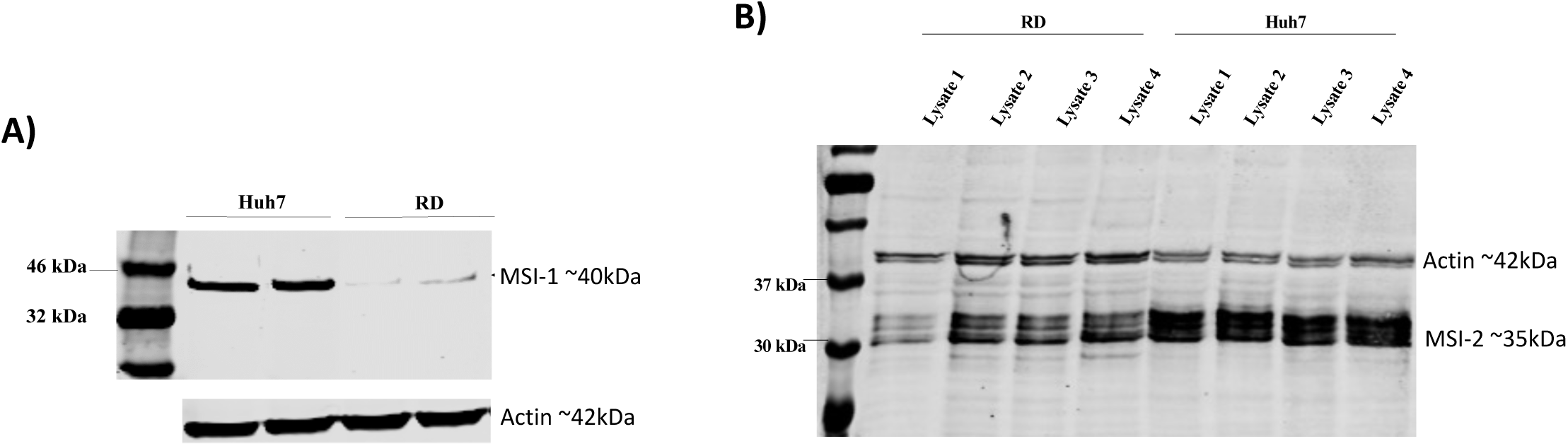
A) MSI-1 and **B**) MSI-2 expression in RD and Huh7 cell lysate analyzed by western blot.

**Supplementary data 7:**
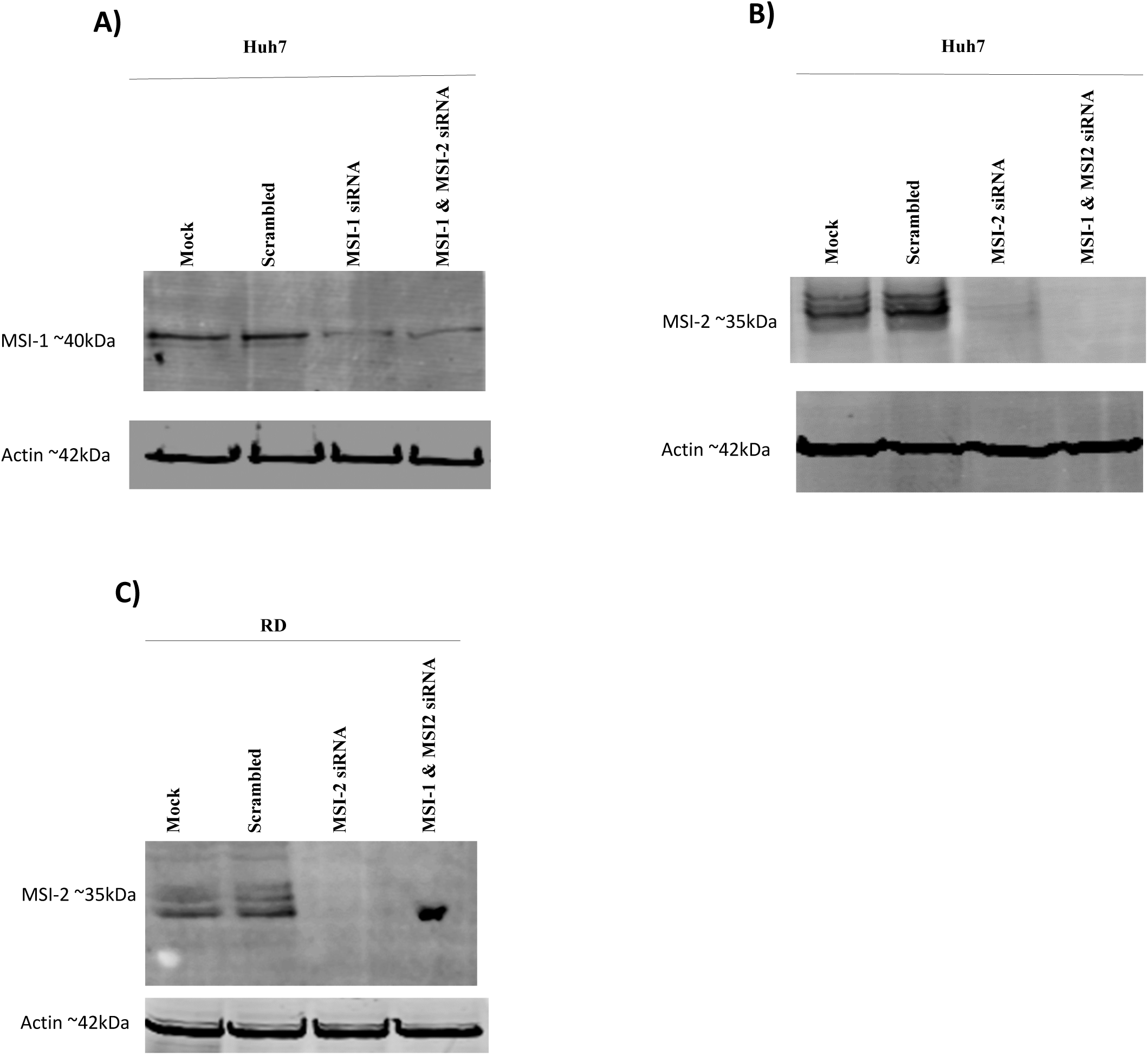
Co-inhibition of MSI-2 and MSI-1 by siRNA in **A)** and **B)** Huh7 cells and **C)** RD cells

## REFERENCES

1. Fong, S.W., Kini, R.M. and Ng, L.F.P. (2018) Mosquito Saliva Reshapes Alphavirus Infection and Immunopathogenesis. Journal of Virology, 92.

2. McPherson, R.L., Abraham, R., Sreekumar, E., Ong, S.E., Cheng, S.J., Baxter, V.K., Kistemaker, H.A.V., Filippov, D.V., Griffin, D.E. and Leung, A.K.L. (2017) ADP-ribosylhydrolase activity of Chikungunya virus macrodomain is critical for virus replication and virulence. P Natl Acad Sci USA, 114, 1666–1671.

3. Petitdemange, C., Wauquier, N. and Vieillard, V. (2015) Control of immunopathology during chikungunya virus infection. Journal of Allergy and Clinical Immunology, 135, 846–855.

4. Ng, L.F. (2017) Immunopathology of Chikungunya virus infection: lessons learned from patients and animal models. Annual review of virology, 4, 413–427.

5. Schneider, M., Narciso-Abraham, M., Hadl, S., McMahon, R., Toepfer, S., Fuchs, U., Hochreiter, R., Bitzer, A., Kosulin, K., Larcher-Senn, J. et al. (2023) Safety and immunogenicity of a single-shot live-attenuated chikungunya vaccine: a double-blind, multicentre, randomised, placebo-controlled, phase 3 trial. Lancet, 401, 2138–2147.

6. Meshram, C.D., Agback, P., Shiliaev, N., Urakova, N., Mobley, J.A., Agback, T., Frolova, E.I. and Frolov, I. (2018) Multiple Host Factors Interact with the Hypervariable Domain of Chikungunya Virus nsP3 and Determine Viral Replication in Cell-Specific Mode. J Virol, 92.

7. Rupp, J.C., Sokoloski, K.J., Gebhart, N.N. and Hardy, R.W. (2015) Alphavirus RNA synthesis and non-structural protein functions. Journal of General Virology, 96, 2483–2500.

8. Tuplin, A. (2015) Diverse roles and interactions of RNA structures during the replication of positive-stranded RNA viruses of humans and animals. J Gen Virol.

9. Frolov, I., Hardy, R. and Rice, C.M. (2001) Cis-acting RNA elements at the 5′ end of Sindbis virus genome RNA regulate minus-and plus-strand RNA synthesis. Rna, 7, 1638–1651.

10. Kendall, C., Khalid, H., Muller, M., Banda, D.H., Kohl, A., Merits, A., Stonehouse, N.J. and Tuplin, A. (2019) Structural and phenotypic analysis of Chikungunya virus RNA replication elements. Nucleic Acids Res, 47, 9296–9312.

11. Troschel, F.M., Minte, A., Ismail, Y.M., Kamal, A., Abdullah, M.S., Ahmed, S.H., Deffner, M., Kemper, B., Kiesel, L., Eich, H.T. et al. (2020) Knockdown of Musashi RNA Binding Proteins Decreases Radioresistance but Enhances Cell Motility and Invasion in Triple-Negative Breast Cancer. Int J Mol Sci, 21.

12. Sutherland, J.M., Siddall, N.A., Hime, G.R. and McLaughlin, E.A. (2015) RNA binding proteins in spermatogenesis: an in depth focus on the Musashi family. Asian journal of andrology, 17, 529–536.

13. Horisawa, K., Imai, T., Okano, H. and Yanagawa, H. (2010) The Musashi family RNA-binding proteins in stem cells. Biomolecular concepts, 1, 59–66.

14. Chavali, P.L., Stojic, L., Meredith, L.W., Joseph, N., Nahorski, M.S., Sanford, T.J., Sweeney, T.R., Krishna, B.A., Hosmillo, M. and Firth, A.E. (2017) Neurodevelopmental protein Musashi-1 interacts with the Zika genome and promotes viral replication. Science, 357, 83–88.

15. Tsetsarkin, K., Higgs, S., McGee, C.E., De Lamballerie, X., Charrel, R.N. and Vanlandingham, D.L. (2006) Infectious clones of Chikungunya virus (La Reunion isolate) for vector competence studies. Vector Borne Zoonotic Dis, 6, 325–337.

16. Pohjala, L., Utt, A., Varjak, M., Lulla, A., Merits, A., Ahola, T. and Tammela, P. (2011) Inhibitors of alphavirus entry and replication identified with a stable Chikungunya replicon cell line and virus-based assays. PloS one, 6, e28923.

17. Lello, L.S., Utt, A., Bartholomeeusen, K., Wang, S., Rausalu, K., Kendall, C., Coppens, S., Fragkoudis, R., Tuplin, A., Alphey, L. et al. (2020) Cross-utilisation of template RNAs by alphavirus replicases. PLoS pathogens, 16, e1008825.

18. Muller, M., Slivinski, N., Todd, E., Khalid, H., Li, R., Karwatka, M., Merits, A., Mankouri, J. and Tuplin, A. (2019) Chikungunya virus requires cellular chloride channels for efficient genome replication. PLoS neglected tropical diseases, 13, e0007703.

19. Bartholomeeusen, K., Utt, A., Coppens, S., Rausalu, K., Vereecken, K., Arien, K.K. and Merits, A. (2018) A Chikungunya Virus trans-Replicase System Reveals the Importance of Delayed Nonstructural Polyprotein Processing for Efficient Replication Complex Formation in Mosquito Cells. J Virol, 92.

20. Clingman, C.C., Deveau, L.M., Hay, S.A., Genga, R.M., Shandilya, S.M., Massi, F. and Ryder, S.P. (2014) Allosteric inhibition of a stem cell RNA-binding protein by an intermediary metabolite. eLife, 3, e02848.

21. Zearfoss, N.R., Deveau, L.M., Clingman, C.C., Schmidt, E., Johnson, E.S., Massi, F. and Ryder, S.P. (2014) A conserved three-nucleotide core motif defines Musashi RNA binding specificity. J Biol Chem, 289, 35530–35541.

22. Arkin, M.R. and Wells, J.A. (2004) Small-molecule inhibitors of protein– protein interactions: progressing towards the dream. Nature reviews Drug discovery, 3, 301–317.

23. Niederhauser, O., Mangold, M., Schubenel, R., Kusznir, E., Schmidt, D. and Hertel, C. (2000) NGF ligand alters NGF signaling via p75NTR and TrkA. Journal of neuroscience research, 61, 263–272.

24. Minuesa, G., Albanese, S.K., Xie, W., Kazansky, Y., Worroll, D., Chow, A., Schurer, A., Park, S.-M., Rotsides, C.Z. and Taggart, J. (2019) Small-molecule targeting of MUSASHI RNA-binding activity in acute myeloid leukemia. Nature communications, 10, 1–15.

25. Lello, L.S., Utt, A., Bartholomeeusen, K., Wang, S., Rausalu, K., Kendall, C., Coppens, S., Fragkoudis, R., Tuplin, A. and Alphey, L. (2020) Cross-utilisation of template RNAs by alphavirus replicases. PLoS pathogens, 16, e1008825.

26. Roberts, G.C., Zothner, C., Remenyi, R., Merits, A., Stonehouse, N.J. and Harris, M. (2017) Evaluation of a range of mammalian and mosquito cell lines for use in Chikungunya virus research. Scientific reports, 7, 1–13.

27. Matkovic, R., Bernard, E., Fontanel, S., Eldin, P., Chazal, N., Hersi, D.H., Merits, A., Peloponese, J.M. and Briant, L. (2019) The Host DHX9 DExH-Box Helicase Is Recruited to Chikungunya Virus Replication Complexes for Optimal Genomic RNA Translation. Journal of Virology, 93.

